# From neural border to migratory stage: A comprehensive single cell roadmap of the timing and regulatory logic driving cranial and vagal neural crest emergence

**DOI:** 10.1101/2022.03.23.485460

**Authors:** Aleksandr Kotov, Mansour Alkobtawi, Subham Seal, Vincent Kappès, Sofia Medina Ruiz, Hugo Arbès, Richard Harland, Leonid Peshkin, Anne H. Monsoro-Burq

## Abstract

Neural crest cells exemplify cellular diversification from a multipotent progenitor population. However, the full sequence of molecular choices orchestrating the emergence of neural crest heterogeneity from the embryonic ectoderm remains elusive. Gene-regulatory-networks (GRN) govern early development and cell specification towards definitive neural crest. Here, we combine ultra-dense single cell transcriptomes with machine-learning and large-scale experimental validation to provide a comprehensive GRN underlying neural crest fate diversification from induction to early migration stages in the frog *Xenopus tropicalis*. During gastrulation, a transient neural border zone state precedes the choice between neural crest and placodes which includes multiple and converging gene programs. Transcription factor connectome and bifurcation analyses demonstrate the early emergence of neural crest fates at the neural plate stage, alongside an unbiased multipotent lineage persisting until after epithelial-mesenchymal transition. We decipher the circuits driving cranial and vagal neural crest formation and provide a broadly applicable strategy for investigating SC transcriptomes in vertebrate GRNs in development, evolution, and disease.

## Introduction

Neural crest cells form a population of multipotent and migratory progenitors found in vertebrate embryos, essential for the peripheral and enteric nervous system, craniofacial structures, endocrine and pigment cells among others. Together with ectodermal placodes, neural crest (NC) cells are evolutionary inventions that support many cell and tissue innovations promoting the vertebrate predatory lifestyle. Shortly after gastrulation, NC cells are induced from the dorsal-lateral “neural border zone” (NB), an ectoderm domain located between the non-neural ectoderm and the neural plate ectoderm (Eames et al., 2020; Plouhinec et al., 2017). In addition to the NC, the NB territory also gives rise to posterior placodes, non-neural ectoderm and the dorsal part of the neural tube (Steventon and Mayor, 2012; Streit and Stern, 1999). Whether these four cell types arise from a common and multipotent early progenitor state, and how fate decisions are orchestrated at the NB during gastrulation remain poorly understood. During neurulation, NC specification and induction progresses as an anterior-to-posterior wave along the edges of the neural plate, with gene programs that define early and immature neural crest cells (e.g. expression of *snail2, foxd3* and *sox8* genes) followed by later pre-migratory programs presaging emigration of NC cells from the NB epithelium as the neural folds elevate and close (e.g. expression of *sox10*, *twist1* and *cdh2 (N-cadherin)* genes) (Bhattacharya et al., 2020; Figueiredo et al., 2017). In addition to this pan-NC program, several regional molecular modules are activated along the anterior-posterior body axis and define subpopulations with specific potential (Ling and Sauka-Spengler, 2019; Tang et al., 2021). How these programs are interconnected with the pan-NC module, and how and when they are activated in pre-migratory NC cells is poorly described. Later, at the end of neurulation, NC cells leave the dorsal ectoderm by a stereotypical epithelium-to-mesenchymal transition (EMT) followed by extensive migration towards a variety of target tissues.

NC biology has been scrutinized during development and evolution, leading to the elucidation of elaborate gene regulatory networks (GRNs) during the last decade (Monsoro-Burq et al., 2005; Simoes-Costa and Bronner, 2016). These networks, however, remain incomplete and do not account for most of the defects observed in human neurocristopathies (Medina-Cuadra and Monsoro-Burq, 2021). This problem is ripe for single cell (SC) transcriptomics, which would enable a full description of NC development over sequential developmental stages, and in comparison to adjacent tissues (e.g. at the neural border) would define the developmental genetic trajectories of the complete NC lineage tree. Most of the recent SC studies on NC cells have mainly explored NC after emigration (Artinger and Monsoro-Burq, 2021; Supplementary File 1 - Table S1). In contrast, pre-migratory NC single cells have received limited exploration, mostly around the EMT stage and on small cell numbers at a specific level of the body axis (Ling and Sauka-Spengler, 2019; Zalc et al., 2021). Earlier on, formation of the NB territory has been defined by expression of a few genes during gastrulation (e.g. *pax3* and *pax7)* (Basch et al., 2006; Monsoro-Burq et al., 2005; Plouhinec et al., 2017), however the timing of NB specification from the rest of the dorsal ectoderm and the circuits driving fate decisions between the four NB-derived cell fates (NC, placodes, non-neural ectoderm and dorsal neural tube) remain to be established (Groves and LaBonne, 2014; Maharana and Schlosser, 2018; Steventon and Mayor, 2012). Furthermore, the timing of lineage decisions in the pre-migratory NC along the anterior-posterior axis, the maintenance of a multipotent NC subpopulation, and the molecular mechanisms driving each state of the pre-migratory NC lineage tree remain unexplored. Here, we used single cell transcriptomes from eight consecutive developmental stages of *Xenopus tropicalis*, featuring 6135 NC cells and 17138 early ectoderm cells, to provide a comprehensive developmental profiling of the NB and the pre-migratory NC. During neurulation, we define several new premigratory NC subpopulations and highlight the transcriptomic trajectories between these early states and eight NC subpopulations emigrating from anterior to vagal levels of the body axis. Interestingly, we find that distinct signatures for prospective NC fates emerge much earlier than previously anticipated, that NC state diversity is maintained upon EMT and that further diversification occurs at the onset of migration. During gastrulation, we explore neural border development and its specification into the NC and placodes. At each stage, we propose a temporal sequence of molecular events underlying these successive transcriptomic states and identify key candidate transcription factors involved in the branching between states. Importantly, we validate several regulatory predictions using transcriptomes and *in vivo* approaches. Last, we propose a model of “dual convergence” for parallel transcriptomic routes driving neural border specification. We therefore provide an extensive gene regulatory network describing the emergence of the neural crest from the ectoderm of vertebrate embryos.

## Results

### Defining the diversity of premigratory neural crest states

Using deeper re-sequencing of single cell (SC) series from whole *X. tropicalis* embryos (Briggs et al., 2018), taken at 8 consecutive developmental stages, followed by updated genome annotation and alignment, we have scrutinized 6135 neural crest (NC) cells from early gastrulation to early migration stages. The larger cell number of our new dataset allowed greater assessment of the cellular diversity in the NC population during early induction (at late gastrulation stages 12-13 and neural plate stages 13-14), during neural fold elevation (stages 16-18), during EMT (neural tube stages 18-20) and at the earliest stages of NC cells emigration (tailbud stage 22).

Through unsupervised Leiden clustering of the whole embryo dataset at each stage (a total of 177250 cells) (Traag et al., 2019), we identified NC clusters by expression of well-established NC genes from induction to migration (gene-supervised approach, Figure 1A-B; Supplementary File 1 - Table S2). From the annotated frog dataset, signatures of NC/non-NC cells were then used to train a classifier which faithfully detected NC cells when applied to a zebrafish whole embryo dataset. This test indicated that the initial gene-supervised analysis had retrieved *bona fide* NC cells (details in Supplementary Materials; Figure 1 – figure supplement 1). Based on their NC-specific expression and low variation across the NC populations, we found that *tfap2b, c9, c3* and *sox8* are highly expressed throughout NC during neurulation, closely followed by *snai2* and *sox10* (Figure 1C). Stage-specific expression variations suggested that the early NC is best labeled by *snai2* at stages 13-16, followed by *tfap2b* from stage 14 to 20, and *c3, c9, sox8* between stages 14 to 18. Collectively, these genes thus define a canonical “pan-NC” signature.

**Figure 1.**
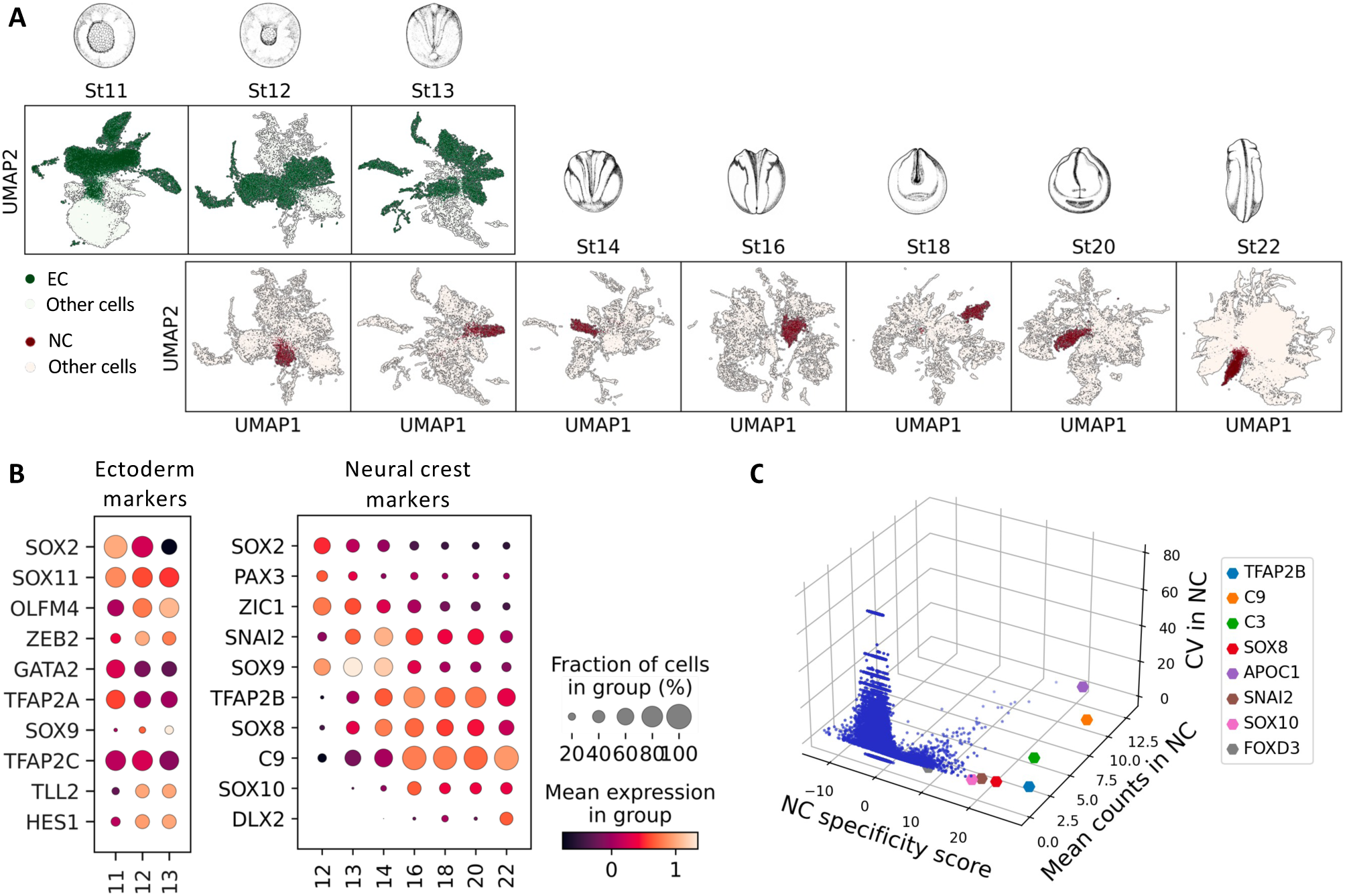
Cell selection for neural crest, ectoderm and neural border. **(A)** Ectoderm (EC, stages 11-13, green) and NC cells (stages 12-22, brown) were selected from a whole embryo SC transcriptome dataset of 177250 cells. **(B)** Dotplots for well-referenced gene expressions used to identify EC and NC at each stage. Dot size represents the number of cells expressing the gene, color represents the average expression level. Neural border was defined as stage 11-13 *tfap2a^+^, zic1^+^* cells. **(C)** 3D scatter-plot of NC score specificity (z-scores), mean gene expression levels (counts) and coefficient of variation (CV) in NC cells, defining a few highly expressed pan-NC genes during neurulation. Additional validation of NC cell selection was done using a binary classifier depicted in Figure 1 - figure supplement 1.

From late gastrulation to emigration stages, reclustering defined sixteen different NC states, compared to 8 described previously (Briggs et al., 2018) (Figure 2A, B). Partition-based graph abstraction analysis (PAGA) (Wolf et al., 2018) showed that most NC clusters are highly interconnected with stronger connectivity between early (1-4), vagal (8, 9, 12 and 13), and cranial (10, 7 and 11) clusters (Figure 2C). Each cluster’s characteristics are described in detail in supplementary text, only new general features are described here (Supplementary file 1, Table S3). First, we identified 10 clusters expressing various levels of *itga4/*integrin-a4, *vim/*vimentin or *fn1*/fibronectin, an explicit mesenchymal NC signature (Figure 2 – figure supplement 1A). These clusters mainly included stage 18 to 22 cells (Figure 2 – figure supplement 1B), stages when the cranialmost NC undergoes EMT and early migration. Using *hox* gene expression, we positioned each cluster along the body axis (cranialmost clusters were *hox-*negative while vagal clusters expressed a range of anterior-to-posterior *hox* genes; Figure 2D). Next, we queried whether all NC cells adopted a similar “stem-like” state upon EMT as proposed recently (Zalc et al., 2021). Instead, we observed high diversity across the mesenchymal clusters: for example, clusters 10, 13, and 15 all undergo a transition to a mesenchymal *vim^+^* state but 10 and 15 expressed the ectomesodermal marker *twi1*, while 13 instead highly expressed *tnc* (Figure 2B, E). Clearly, in the large dataset considered, the diversity of states was maintained as cells transitioned from pre-EMT stages to EMT and early migration. Among the mesenchymal clusters, cluster 16 was the only one enriched simultaneously for late pan-NC markers (*tfap2b, sox10, c9*) and differentiation markers, muscle genes *actc1* and *myl;* (Figure 2E, Figure 2 – figure supplement 2). This NC dataset does reveal other cranial or vagal cluster expressing determination or differentiation markers. This observation matches recent lineage tracing in cranial NC and supports that most NC cell fates are determined post-EMT (Baggiolini et al., 2015; Morrison et al., 2021).

**Figure 2.**
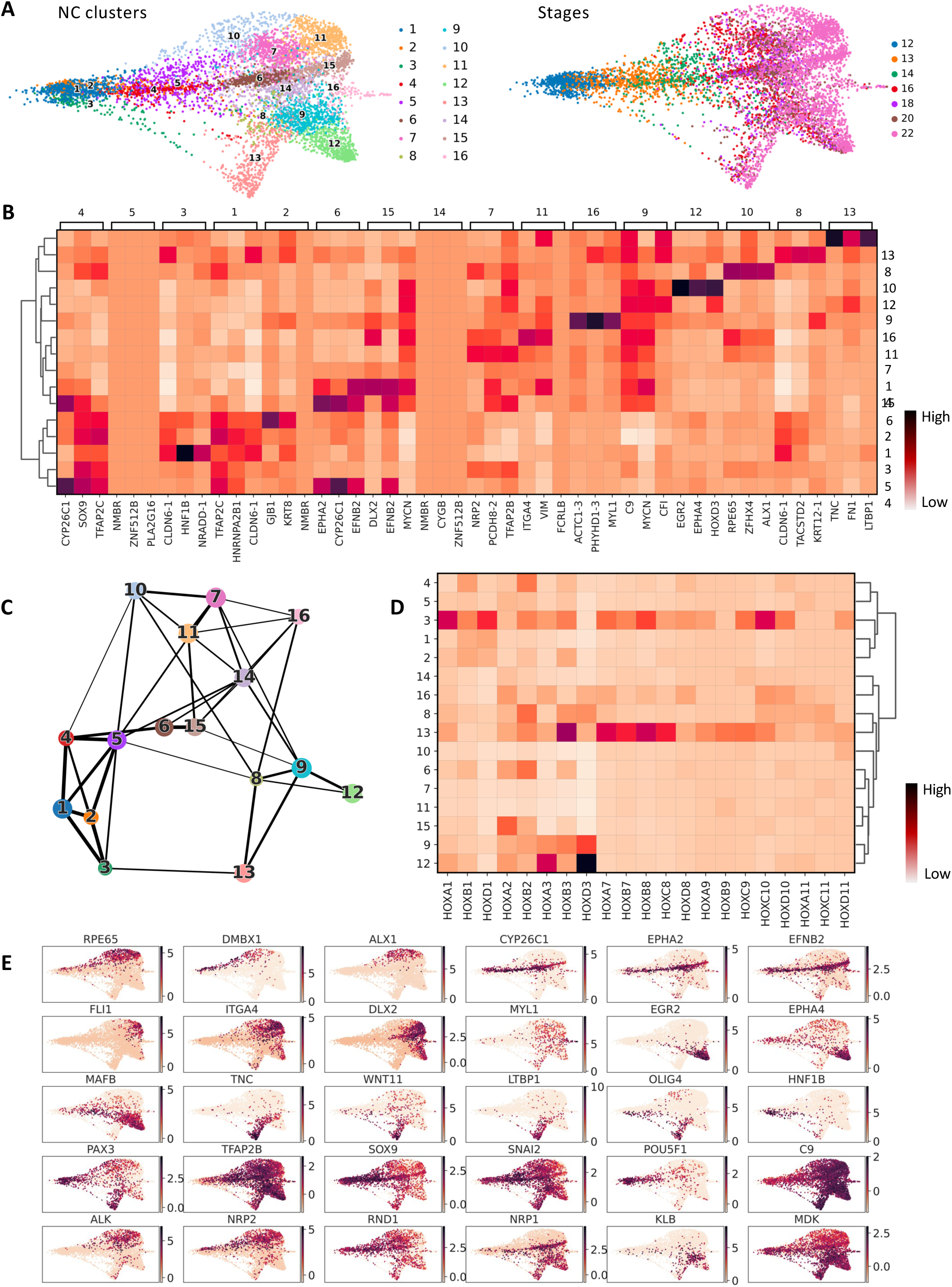
Premigratory neural crest transcriptome heterogeneity. **(A)** Leiden clustering revealed 16 distinct states (clusters) before and during EMT (developmental stages 12-22). 1- Early unbiased / tfap2c^+^; 2- Early unbiased / gjb1^+^; 3- Premigratory sacral / hnf1b^+^; 4- Premigratory early R4 /cyp26c1^+^; 5- Premigratory unbiased; 6- Premigratory late R4 / cyp26c1^+^; 7- Migratory bipotent cranial / nrp2^+^; 8- Migratory rhombencephalic / hoxb2^+^; 9- Migratory bipotent vagal / hoxd3^+^; 10- Premigratory early cranial / alx1^+^; 11- Migratory cranial / itga4^+^; 12- EMT Cardiac / egr2^+^; 13- Migratory ENSp / tnc^+^; 14- EMT unbiased; 15- Migratory late R4 / efnb2^+^; 16- Migratory muscle-like /myl1^+^ **(B)** Top-3 enriched genes for each cluster (top and bottom lines), with their expression in the other clusters and hierarchical clustering between cluster. **(C)** PAGA estimates cluster connectivity where line thickness increases with stronger connections**. (D)** Hox gene signature of each cluster used to approximate their position along the antero-posterior body axis. **(E)** Expression of key cluster-specific genes, including *rpe65, dmbx1, alx1* (cluster 10), *cyp26c1, epha2, efnb2* (clusters 4/6/15), *fli1, itga4, dlx2* (cluster 11), *egr2, epha4, mafb* (cluster 12), *tnc, wnt11, ltbp1* (cluster 13), early *olig4, hnf1b* (cluster 3) and muscle-like NC specific *myl1* (cluster 16). Genes expressed broadly in NC cells define a “canonical NC” signature: early *pax3, tfap2b, sox9, snai2, and c9.* Multipotency-related genes are present mostly until mid-neurula stage *(pou5f1). Alk, nrp2, rnd1* and *nrp1, klb, mdk* are early specifiers of cranial and vagal NC respectively. Detailed cluster characteristics can be found in Supplementary File Table S3 and Text, and in Figure 2 - figure supplements 1-5.

Next, we were able to define multiple progenitor states linked to the later clusters defined above, and consequently delineate transcriptional dynamics of cranial, vagal and enteric nervous system progenitors between late gastrulation and EMT. Late cranial clusters 10 (*nrp2*^+^*rpe65*^+^*hox*^-^) and 11 (*nrp2*^+^*dlx2*^+^*hox*^-^) were related to cluster 7 (*nrp2*^+^ *hox*^-^) while late vagal clusters 12 (*mafb*^+^*epha4*^+^) and 13 (*tnc*^+^*wnt11*^+^*ltbp1*^+^) were linked to cluster 9 (*nrp1*^+^). Cluster 13 was also linked to the early cluster 3 (*hnf1b^+^*). In turn, clusters 7 and 9 were related to the unbiased cluster 5 (*tfap2c*^+^*sox9*^+^). The *nrp1*^+^ late cranial cluster 15 was linked to clusters 6 and 4, all of which specifically expressed *cyp26c1, epha2 and efnb2.* Cluster 16 (*myl1*^+^, Figure 2 - figure supplement 2) was related to unbiased cluster 14, which was in turn linked to unbiased clusters 5, 1 (*c9^+^sox9^+^*) and 2 (*gjb1*^+^) (Figure 2 – figure supplements 3-5; Figure 3).

**Figure 3.**
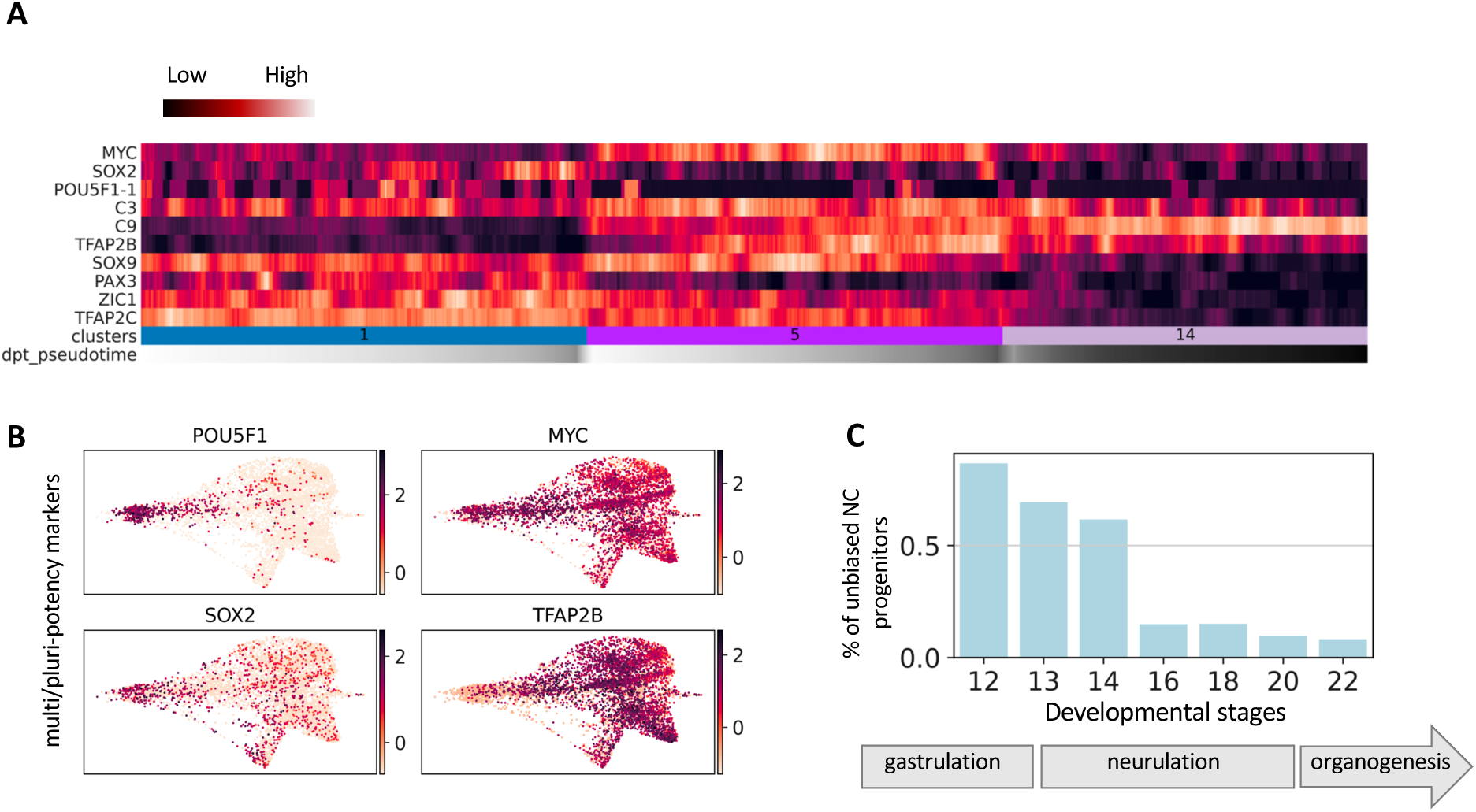
Characteristics of the unbiased NC clusters. **(A)** Expression of pluripotency and “stem” neural crest genes (each bar is a group of 20 cells) in unbiased clusters 1, 5, 14. Pseudotime represents how far the cell has advanced along a given transcriptional path. **(B)** UMAP plots depicting expression of pluripotency-related markers. **(C)** Proportions of the unbiased cells at each developmental stage of the NC dataset.

Interestingly, most clusters gathered cells from multiple stages, indicating that a given transcriptome state can be reached by different cells over long developmental periods (Figure 2 – figure supplement 1B). For example, cranialmost cells of cluster 10 are generated over the entire duration of neurulation from neural plate stage 13 to migration stage 22. This reflects how NC cells of a similar state can be continuously generated over the course of development at a given portion of the neuraxis. In contrast, cranial clusters 11 and 15, two migratory clusters highly expressing *vimentin (vim^+^),* are suddenly generated at post-EMT stage 22. This could reflect an abrupt step of mesenchymalisation in the epithelial-to-mesenchymal transition process of cranial NC, reflected in their transcriptomes.

Moreover, we detected cells with an immature pan-NC signature throughout neurulation and EMT, representing a long-lasting potentially “multipotent” and “stem-like” population (1, 5, 14; Figure 3). These unbiaised progenitors formed 72% of all NC cells during induction (gastrula and neural plate stages 12-14), 15% at pre-migratory and EMT stages 16-18, and 9% among tailbud stage 22 NC cells (Figure 3C). This dataset thus identifies the stem-like and partially biased premigratory NC populations; details the temporal dynamics of multipotency genes and the sequential appearence of regional modules; and characterizes the progressive activation of mesenchymal markers across a diversity of cell states. In sum, we uncover the highly diverse transcriptomes underlying premigratory NC cell biology along a large, cranial to vagal segment of the body axis.

### Connectome analysis by intersecting sc-transcriptomics, morpholino RNA-seq and ChIP-seq

To expand the network of genes involved in the NC-GRN, we used an input list of 1417 transcription factors (TFs) (Blitz et al., 2017) and applied GRNboost2 on the NC dataset to link transcription factors to their potential targets in the NC transcriptome. We retrieved a network of 16978 potential TF-targets with a median of 22 connections per TF. Among the most connected genes for each stage (Table 1), we retrieved the previously known neural border specifiers *pax3* and *zic1* at NC induction stages, as well as *zic3, olig4, sox9*, and the anterior NC marker *dmbx1*. At pre-migratory stages, the prominent nodes included the NC specifiers *tfap2b, sox10 and snai2*, the anterior NC markers *rpe65* and *alx1*, and the hindbrain hox gene *hoxb3*. At migration stages, *tfap2e*, *mycn, dlx2* and *egr2* (*krox20)* displayed both high expression and many connections.

**Table 1.** Highly connected genes in the neural crest network predicted by GRNboost2.

| Gene | Degree centrality | Betweenness centrality | Max. expression stage | NC specificity |
| --- | --- | --- | --- | --- |
| TFAP2C | 28.028933 | 21.510673 | 12 | 6.59 |
| ZIC3 | 16.455696 | 7.885226 | 12 | 3.73 |
| ZIC1 | 8.137432 | 2.351524 | 12 | 9.34 |
| ZIC2 | 7.233273 | 2.026896 | 12 | 3.62 |
| DMBX1 | 3.616637 | 11.876445 | 12 | 1.57 |
| PAX3 | 3.435805 | 2.211830 | 12 | 5.93 |
| SOX9 | 11.573237 | 8.575830 | 13 | 14.07 |
| OLIG4 | 4.882459 | 2.642370 | 13 | 4.54 |
| MAFB | 7.956600 | 4.174946 | 14 | 09.08 |
| SNAI2 | 5.244123 | 0.927599 | 14 | 20.03 |
| SOX8 | 4.520796 | 2.071516 | 14 | 20.60 |
| RPE65 | 1.446655 | 0.534404 | 14 | 6.49 |
| TFAP2B | 8.499096 | 4.135991 | 16 | 25.47 |
| HOXB3 | 3.978300 | 0.959282 | 16 | 2.35 |
| SOX10 | 3.797468 | 0.705758 | 16 | 15.40 |
| NR6A1 | 3.797468 | 0.706433 | 16 | 9.18 |
| ZFHX4 | 2.169982 | 0.791908 | 16 | 4.83 |
| ALX1 | 2.169982 | 1.627017 | 16 | 3.69 |
| EPHA2 | 1.989150 | 1.403078 | 18 | 3.77 |
| E2F3 | 1.446655 | 2.256467 | 18 | 0.68 |
| TFAP2E | 5.786618 | 1.850976 | 20 | 15.21 |
| EGR2 | 5.424955 | 2.505897 | 20 | 5.96 |
| MYCN | 17.359855 | 12.737248 | 22 | 8.81 |
| DLX2 | 12.839060 | 8.413243 | 22 | 6.88 |
| HOXD3 | 8.137432 | 3.417475 | 22 | 4.96 |
| EEF1D | 4.701627 | 0.795235 | 22 | 9.55 |
| ZBTB16 | 3.074141 | 1.013930 | 22 | 2.80 |
| TFEC | 2.169982 | 0.562422 | 22 | 0.10 |
| VIM | 2.169982 | 0.879581 | 22 | 2.43 |

In order to experimentally test the predicted network, we sequenced NB/NC microdissected ectoderm after *in vivo* knockdown of selected nodes *pax3* and *tfap2e,* using previously validated antisense morpholino oligonucleotides (MO) (Figure 4A, Hong et al., 2014; Monsoro-Burq et al., 2005). Expression of 1333 transcripts were decreased in Pax3 morphant NB, confirming that Pax3 is essential to activate a large NB/NC gene set (Figure 4A, Supplementary file 1 – Table S4). We verified direct Pax3-binding targets *in vivo* using ChIP-seq on mid-neurula stage embryos (Figure 4B) and identified 657 potential targets in the whole embryo, of which 475 were expressed in the SC NC dataset (Supplementary file 1 – Table S6), including known direct targets, *e.g. cxcr4* and *prtg* (Plouhinec et al., 2014; Xu et al., 2018). Moreover, 80 of the ChIPseq-validated targets were predicted by GRN-Boost2 modelling including *psmd4, psen2, sp7, notch1, hnf1b.* In sum, we provide here a genome-wide Pax3 NC-GRN with three complementary approaches: co-expression predictions, Pax3 depletion and Pax3 chromatin-binding (Figure 4C). Using a similar approach, we confirmed TFAP2e as an important regulator at early EMT stages. Among 848 targets of TFAP2e predicted from scRNA-seq, 99 showed changed expression after TFAP2e knockdown in NC *in vivo* (e.g., *tfap2b, sox10, sncaip;* Figure 4 – figure supplements 1-2; Supplementary file 1 – Table S5). Moreover, using ChIP-seq for TFAP2e, we identified 642 targets expressed in the NC dataset, among 805 targets for the whole embryo, including top-scored *rmb20, pim1, arl5b* and *tfap2a* (Supplementary file 1 – Table S7). In both Pax3 and Tfap2e cases, we find that the three approaches result in partially overlapping gene target lists due to their use of different parameters (stage-wise, expression-wise, etc; Figure 4C, see Materials and Methods). Together, these data provide an enlarged and validated NC-specific genome-wide connectome for two key NB/NC specifiers.

**Figure 4.**
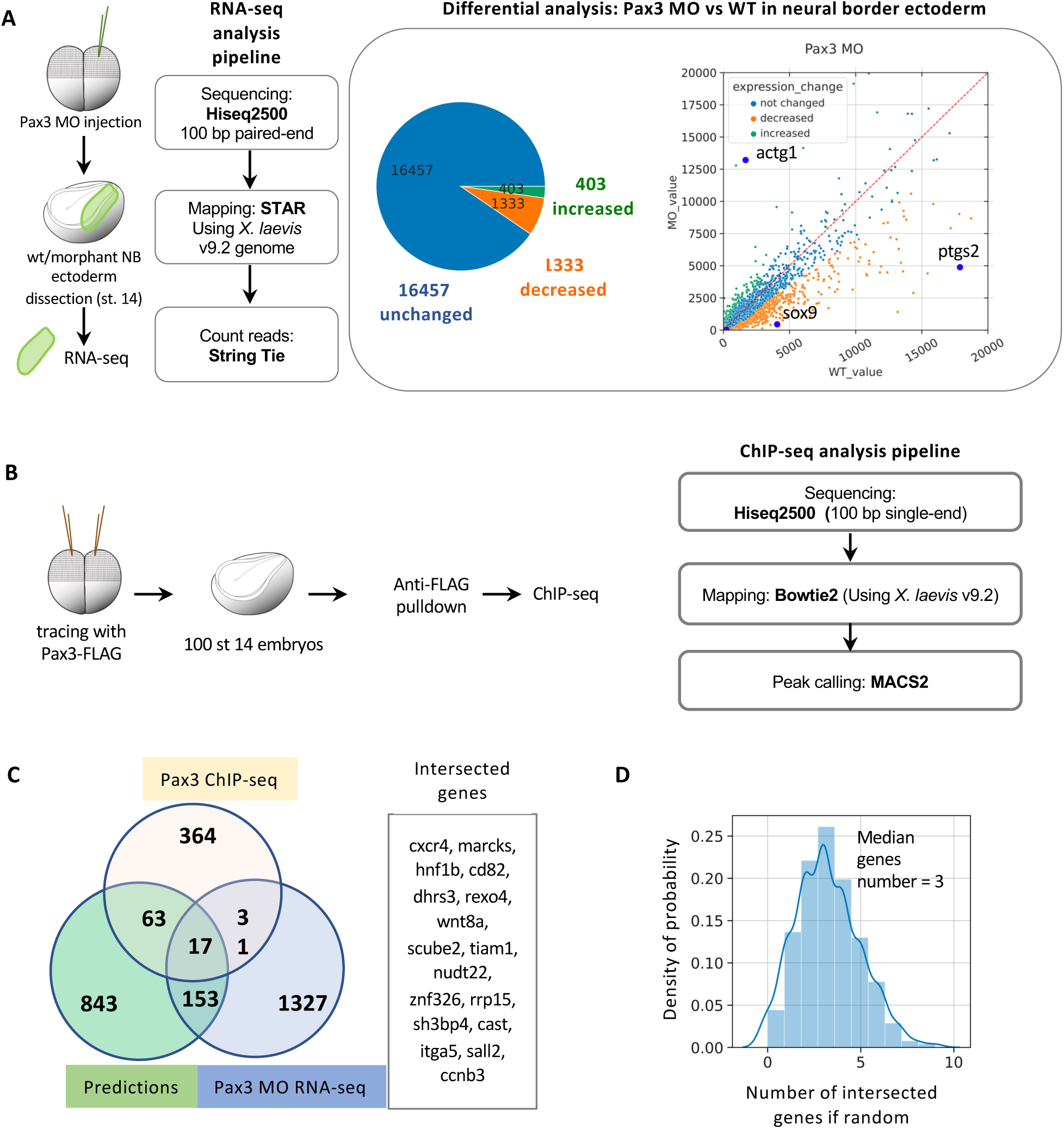
Pax3 and TFAP2e connectome generation and validation. (A) RNA-sequencing was done on microdissected NB ectoderm (stage 14), either wild-type (wt) or following Pax3 depletion in vivo (Pax3 MO). Using standard differential analysis, 1333 genes were found downregulated after Pax3 depletion, including sox9, olig4, hnf1b, msx1 (Table S4). (B) Tracing amounts of Pax3-FLAG expression were used to pull-down Pax3-bound chromatin in vivo without phenotypic modification, followed by standard ChIP-seq analysis pipeline, identifying 657 direct target genes among which 475 were expressed in the NC dataset. (C) Venn diagram compares Pax3 target genes validated by ChIP-seq, MO-RNA-seq and GRN-boost2 modeling. The 17 genes linked to Pax3 by all three methods are listed here (full lists in Supplementary file 1 – Tables S4 and S6). (D) Random sampling (bootstrap) was used to test the average number of genes present in the intersection by the three methods if random chance was applied to these datasets. At random, an average of 3 genes would be found in the intersection compared to 17 found here (C). Similar approach was applied to TFAP2e in Figure 4 - figure supplement 1, 2 and Table S5.

### Branching towards biased premigratory neural crest subpopulations is controlled by key transcription factors

To explore the temporal dynamics of TF expression that may specify decision points in the development of pre-migratory NC, we used tree inference (ElPiGraph) and advanced pseudotime downstream analysis focused on fate biasing using scFates (Albergante et al., 2020). We thus explored potential branch-specific transcriptional regulation from the calculated pseudotime, in order to determine not only the gene-to-gene dependency but also the temporal order in which their functions may be accomplished. ElPiGraph approximates datasets with complex topologies to build the graph structure, with the limitation that it cannot be applied to large datasets with many potential branches. Thus, using the principal graph constructed with PAGA (Figure 2B), we sub-selected cells around three main bifurcation points in the NC lineage tree and then applied scFates branching analysis (Figure 5A-F).

**Figure 5.**
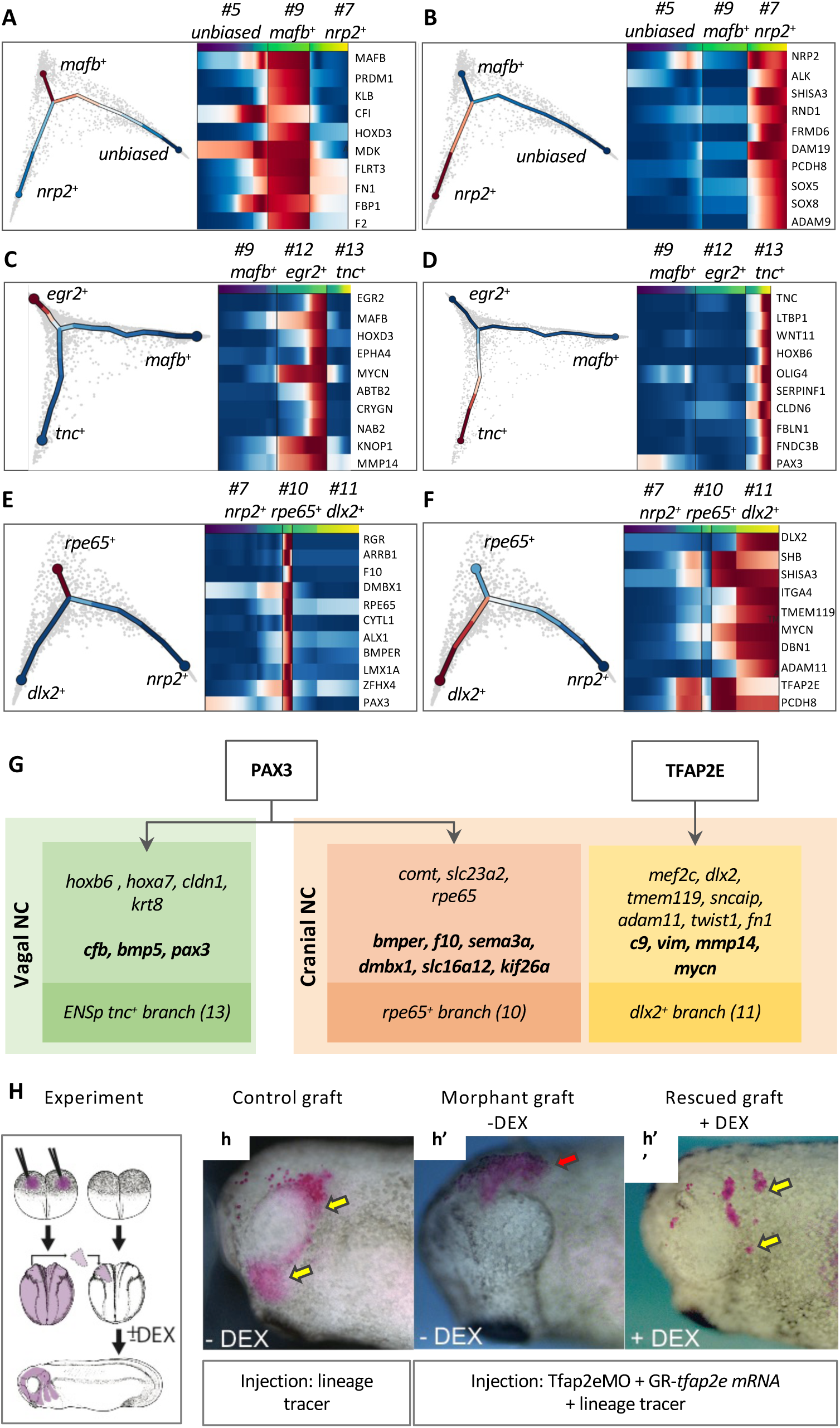
NC branching analysis. Transcriptomes of cells subselected around a chosen bifurcation point were analyzed using tree inference and pseudotime downstream analysis, yielding gene programs accompanying each trajectory. Gene programs for bifurcation of (A-B) premigratory unbiased cluster 5 into cluster 9 (A) and cluster 7 (B); (C-D) of migratory bipotent vagal cluster 9 into clusters 12 (C) and 13 (D); (E-F) of migratory bipotent cranial cluster 7 into clusters 10 (E) and 11 (F). (G) MO-mediated depletion and ChIP-seq was used to test Pax3 function in either vagal branching (cluster 12 vs 13) or cranial branching (cluster 11 vs 10) in vivo. Branch-specific genes with modified expression after Pax3 depletion (standard letters) or bound by Pax3 (in bold letters) are indicated. Using a similar approach, TFAP2e function in cranial branching (cluster 10 vs 11) was validated in vivo. (H) To test TFAP2e function in cranial NC migration, premigratory NC (traced in pink) was grafted into wild-type host embryos. In comparison to control (wt) NC active migration (h, yellow arows), TFAP2e morphant NC cells do not emigrate towards craniofacial areas and remain at the graft site (h’, red arrow). In contrast, reactivation of TFAP2e-GR in morphant cells upon EMT stage restores NC cell migration (h”, yellow arrows).

### Cranial versus vagal bifurcation at the end of neural plate stage

Cranial NC cells emerge from the neural tube anterior to the otic vesicle while vagal NC cells form from the hindbrain region adjacent to somites 1–7 (Le Douarin and Kalcheim, 1999). Our data indicated that the first cells biased towards vagal or cranial populations arose from the unbiased cluster 5 around early neural fold stage 14. Although we did not see a separation of cluster 5 into two populations at the chosen level of clustering, we still observe an early internal predisposition marked by the expression of early cranial (*nrp2)* and vagal (*mafb*) markers in sub-regions of cluster 5: cluster 5 cells that highly express *nrp2* were 7 times closer to the cranial state, while cells that highly express *mafb* were 1.5 times closer to the vagal state (Figure 5 – figure supplement 1A and B). Through branching analysis, we uncovered gene programs governing the bifurcation, consisting of ‘early’ genes activated before bifurcation and ‘late’ genes with continued expression in each branch (late): *nrp2*, *alk*, *rnd1*, *adam19* (early) and *shisa3*, *frdm6* (late) for the cranial branch, and *mafb*, *klb*, *mdk* (early) and *prdm1*, *cf1*, *hoxd3* (late) for the vagal branch (Figure 5A-B). By high specificity and early expression, *nrp2 and mafb* were the best early predictors of branching between the cranial and vagal populations (Figure 2E). These results match *in vivo* analyses on expression of key NC regulators, such as Mafb during cardiac NC specification or Alk in cranial NC migration (Gonzalez Malagon and Liu, 2018; Tani-Matsuhana and Inoue, 2021).

### Vagal to enteric split and cranial subdivisions at neural fold stages

From the apparently homogeneous clusters 7 and 9, branching analysis predicted candidates regulating subsequent bifurcation. For the cardiac NC cluster 12, early-enriched transcripts were *mafb, mycn, prdm1, nolc1* and *eef1d,* late ones being *egr2, hoxd3, epha4* and *abtb2*. For the ENSp cluster 13, early markers were *olig4* and *fbn2*. Due to its significantly increased expression in later ENSp cells, *pax3* was identified as a late actor, but the expression-pseudotime heatmap showed that *pax3* was already expressed prior to branching in ENSp progenitors and increased afterward (Figures 2D and 5D). Therefore, we also assigned *pax3* to the early ENSp branch. Later ENSp gene program consists of *tnc, ltbp1, wnt11,* and *hoxb6*. In order to test experimentally if ‘early’ factors affected expression of ‘late’ genes, we examined which late ENSp genes may be targets of the early TF Pax3, through ChIP-seq and depletion analyses. Some genes were bound by Pax3 (*cfb, bmp5),* while others showed increased (*krt8)* or decreased (*cldn1, hoxb6*, hoxa7) expression without evidence for Pax3 binding (Figure 5G, Figure 5 – figure supplement 1C).

Similar analysis on cranial NC clusters 7-10-11 showed that branch 10-specific genes were *rpe65, dmbx1, rgr, lmx1a,* and *zfhx4*, while late gene programs included *alx1* and *bmper* (Figure 5E-F). Interestingly, *pax3* was expressed early in the cranial bifurcation from 7 towards 10 and 11; expressed in cluster 7 cells, *pax3* was specifically enriched in *rpe65^+^* cluster 10. The Pax3 ChIP-seq and MO datasets revealed that *rpe65^+^* branch-specific transcripts *comt, slc23a2* and *rpe65* were decreased after *pax3* depletion while others, *bmper, f10, sema3a, dmbx1, slc16a12* and *kif26a,* were bound by Pax3 *in vivo* (Figure 5G). On the other hand, *tfap2e* expression initiated before bifurcation and was enriched in cluster 11 relative to cluster 10. Using TFAP2e depletion and ChIP-seq, we also validated that TFAP2e depletion reduced the expression of nine *hox^-^dlx2^+^*branch-specific genes (e.g. early gene *mef2c*, and late genes *dlx2, mmp14, vim*), and that TFAP2e bound four other genes in cluster 11 signature (*c9, vim, mmp14 and mycn,* Figure 5G, Figure 5 – figure supplement 1C). While TFAP2e is essential for NC induction (Hong et al., 2014), direct regulation of *vim* and *mmp14* suggested that TFAP2e may also control cranial NC EMT and migration. We tested this later role *in vivo* using low-level depletion of TFAP2e which still allowed initial NC induction. Specifically, a pre-EMT NC explant co-injected with low-levels of TFAP2e MO, mRNA encoding a dexamethasone-inducible TFAP2e and a lineage tracer, was grafted into a wild-type control embryo prior to EMT (stage 17, Figure 5H). Morphant NC remained at the grafted site while wild-type cells efficiently populated the craniofacial areas (Figures 5Hh, h’). Importantly, when TFAP2e was re-activated in morphant NC at EMT stage, cell migration was restored and lineage-traced cells were found along NC cranial migration routes (Figure 5Hh”). Together, these results validate branching analysis predictions and demonstrated that TFAP2e regulates expression of EMT effectors and cranial NC migration *in vivo*.

In conclusion, using computational approaches we have defined and analyzed three main bifurcation points in the premigratory NC dataset: from unbiased to vagal and cranial NC, from vagal NC to cardiac and ENSp fates, and from early cranial to either *rpe65^+^* or *dlx2^+^*cranial NC. For each branch, we defined specific gene programs, including early actors predicted to trigger specific states. Lastly, we validated the link between numerous branch-specific genes and two of the early regulators, Pax3 and TFAP2e.

### Coexisting neural border, ventral and dorsal ectoderm gene programs specify neural crest and placodes

#### Neural border cell signature displays enriched ventral and dorsal gene expressions which are regulated by Pax3

To unveil the molecular mechanisms that distinguish NC and placode induction at the neural-nonneural ectoderm border zone during gastrulation, we collected data for all ectoderm cells from stages 11-13 and identified 10 cell clusters (Figures 1A and 6A). The NB zone is an ectodermal area located in-between *sox2^+^* neuroepithelium (NE) and *epidermal keratin^+^* nonneural ectoderm (NNE), co-expresses *tfap2a, zic1* and *pax3* and gives rise to both NC and placodes (Figure 6A, B and Figure 6 - figure supplement 1B) (Seal and Monsoro-Burq, 2020). The early developmental dynamics of this ectodermal area has not yet been described at the single-cell level, and it remains unknown if NB cells resemble adjacent progenitors or if they exhibit a specific gene signature. On the force-directed graph plot, *tfap2a^+^zic1^+^* cells (zone 3) did not appear as an individual cluster, rather as cells spread out amongst clusters 1, 2, 5 and 6 (Figure 6A, Figure 6 – figure supplement 1A). While we observed low cell density specifically for the NB zone between stages 11 and 12, we excluded a stage-related sampling issue since NE/NNE areas presented normal density. Rather this could relate to a faster transcriptional transition of NB cells compared to NNE cells: if early ectoderm cells transit through the NB state quickly before switching towards NC or placode states, fewer cells would be captured. Alternatively, such a plot could be obtained if the NB zone was highly heterogeneous and contained a mosaic of states (NB, NNE or NE).

**Figure 6.**
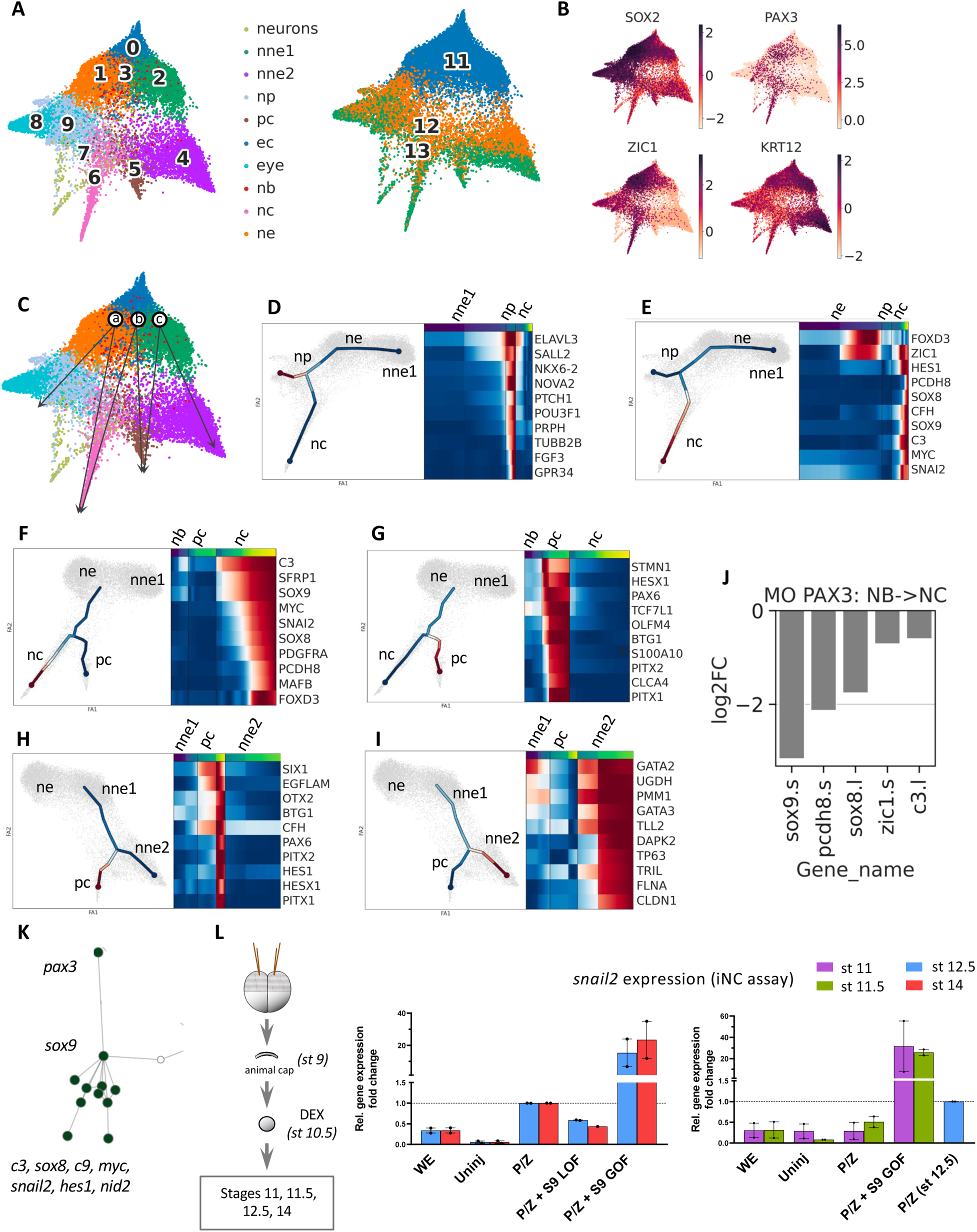
Ectoderm branching and validation of Sox9 early role in NC induction. (A) Forced-directed graphs for the Ectoderm dataset, with cluster numbers and gastrula stages indicated. (B) UMAP plots for genes marking the 3 major ectoderm areas: sox2 is enriched in the nascent neural ectoderm while krt12 (epidermal keratin 12) is highest in the non-neural ectoderm. Unlike the other areas, the NB zone does not display a specific gene signature and is usually depicted by pax3 expression or the overlapping expressions of ventral (tfap2a) and dorsal (zic1) genes (red cells in A, located across dorsal and ventral clusters). (C) Based on the forced-directed graph we hypothesized several paths for NC and PC development at stages 11-13, considering two possible origins for both populations (NP and NB for NC, and NNE and NB for PC). (D) Early NNE (NNE1)->NP gene program, (E) NE->NC gene program, (F) NB->NC gene program, (G) NB->PC gene program, (H) NNE1->PC gene program, (I) NNE1->later NNE (NNE2) gene program (J) Connectome and scFates branching analysis revealed that Pax3 was an important node in the predicted GRN and an early gene for branch NB->NC. In vivo, Pax3 depletion impacted expression of several NB->NC branch-specific genes, including sox9, c3 and foxd3. (K) Analysis of the Ectoderm connectome suggested a novel epistasis relationship between pax3, sox9 and other downstream NC specifiers. (L) In iNC assay, Sox9 acted downstream of Pax3 and was essential for activating the downstream NC program. Additionally, at gastrula stages, a time point at which pax3/zic1 activation does not yet induce snail2 expression in iNC, activating Sox9 was sufficient to obtain high levels of snail2 precociously. RT-qPCR analysis showing relative snail2 expression fold change in iNC at late gastrula and early neurula stages. WE - whole embryo; Uninj - uninjected animal caps; P/Z - pax3-GR + zic1-GR iNC; S9 LOF - sox9 loss-of-function (LOF) or gain-of-function (GOF).

We then identified a detailed signature for the *tfap2a^+^zic1^+^*cells (zone 3). Several transcripts were enriched from stage 11: *tfap2c, pax3, sox9, hes1, gmnn* and *myc* (Figure 6 – figure supplement 1D). Most of these transcripts also formed main nodes in the whole Ectoderm connectome (10085 gene connections, Table 2). We examined whether Pax3, previously shown to appear as early as stage 10.5 *in vivo* (de Crozé et al., 2011), triggers expression of NB zone signature genes *in vivo*; in Pax3 morphant NB transcriptome, we found decreased expression of the other NB genes *sox9, axin2, zic3* and *zic1,* while early NE and NNE marker *lhx5.l* was increased (Figure 6 – figure supplement 1E). These results verify and consolidate Pax3 as a major activator for the early NB signature.

**Table 2.** Highly connected genes in the ectoderm network predicted by GRNboost2.

| Gene | Degree centrality | Betweenness centrality | Mean nc/pc/nb specificity | NC specificity | PC specificity | NB specificity |
| --- | --- | --- | --- | --- | --- | --- |
| TFAP2C | 70.319635 | 6.203872 | 8.778883 | 17.587187 | 3.636364 | 5.113099 |
| TFAP2A | 67.123288 | 5.770185 | 11.238043 | 4.836522 | -0.884711 | 29.762320 |
| ZIC1 | 47.488584 | 1.674848 | 15.645578 | 18.120411 | -8.226690 | 37.043015 |
| HES1 | 30.136986 | 0.815641 | 13.273738 | 27.994427 | 11.076355 | 0.750431 |
| SOX9 | 24.657534 | 0.463349 | 11.848502 | 38.367329 | -4.275626 | 1.453803 |
| MYC | 17.808219 | 0.340629 | 9.269785 | 22.733620 | 4.699314 | 0.376421 |
| SNAI2 | 17.579909 | 0.192783 | 7.027262 | 27.004408 | -3.017278 | -2.905344 |
| PITX1 | 16.438356 | 0.115279 | 5.463268 | -3.656815 | 21.928877 | -1.882259 |
| PAX3 | 14.611872 | 0.172905 | 5.509155 | 15.614752 | -3.379381 | 4.292094 |
| SOX8 | 12.100457 | 0.094409 | 5.176289 | 19.021635 | -2.535636 | -0.957131 |
| C3 | 9.817352 | 0.019518 | 12.694010 | 36.559040 | 6.013386 | -4.490397 |

#### The NB zone contributes to NC and PC in parallel to convergent contributions from neural plate and non-neural ectoderm progenitors

Based on the force-directed graph, we hypothesized three possible developmental routes leading to NC and placodes from stage 11 ectoderm cells: a) NE → NC; b) NB zone → PC and NC; c) NNE → PC (Figure 6C); these possibilities are consistent with current models of NC and PC formation, the “neural border zone model” (route b) and the “neural vs non-neural” model (routes a & c) (Figure 6C) (Maharana and Schlosser, 2018; Roellig et al., 2017; Seal and Monsoro-Burq, 2020). Interestingly using branching analysis, no direct route was found between early NNE (nne1) and NC, or between NE and PC, confirming biological features, such as the partial neuralization needed for NC induction or the close relationship between placodes and NNE (Alkobtawi et al., 2021; Briggs et al., 2018). Since ElPiGraph cannot be applied on the whole dataset, we applied branching analysis to each potential route and compared the resulting gene programs underlying NC vs PC fate decisions. For route (a), branching towards NP from the NE state involved early genes *elav3 and sall2* and late genes *nkx6* and *tubb2b*, consistent with current knowledge on neural plate induction (*sall2, nkx6*) and primary neurogenesis (*elav3, tubb2b*) (Figure 6D, Exner et al., 2017). Route (a) branching towards NC involved early genes *zic1* and *foxd3* and late genes *c3, myc, sox8, sox9, pcdh8* and *snai2* (Figure 6E). In comparison, NC cells emerging from the NB zone by route (b) expressed *c3* and *sox9* before bifurcation, while NC markers *foxd3*, *sox8, pcdh8, snail2* were enriched after splitting (Figure 6F). Similarly, we explored both proposed routes of placodal development: route (b) from NB zone showed early enrichment of *tcf7l1, hesx1* and late for *stmn1, pax6, pitx1/2* (Figure 6G) while route (c) from NNE exhibited early enrichment of placode specifiers *six1, otx2*, and late expression of placode markers *egflam, pax6, pitx1/2*, Figure 6H). Last, the route (c) branching for NNE formation confirmed known developmental dynamics with early enrichment for *gata2* and late for ectoderm stem cell marker *tp63* and epithelial cells *cldn1* (Figure 6I) (Haas et al., 2019). This hierarchy of gene expression, some likely to respond to external signals as well, along the different branches thus opens avenues to further elaborate each of the NC- and PC-GRNs.

#### Different gene programs can lead distinct progenitors towards a similar state

Interestingly, according to the route studied, we found that (i) distinct genes were activated to obtain the same state, and that (ii) some genes were activated with different expression dynamics relative to different bifurcations. For example, during the NB→NC transition (route b, Figure 6F), *sox9* and *c3* were activated early (before bifurcation) suggesting that they could play a part in the fate decision network from NB progenitors. In contrast, during the NE→NC gene program (Figure 6E) *sox9* and *c3* were late genes while *foxd3* and *zic1* were expressed early. This observation suggested a new model of fate decisions in the developing ectoderm, where parallel and distinct genetic programs activated in distinct ectoderm progenitors may lead to a similar state.

Last, NB zone-specific gene *pax3* was expressed prior to bifurcation in the route (b), NB→NC gene program, and activated expression of late NC branch markers *sox9, sox8, zic1, pcdh8* and *c3* (Figure 6J). Moreover, the Ectoderm connectome described NC genes connected to the rest of the network through Pax3 and Sox9 (Figure 6K), suggesting that Sox9 might play a yet undescribed function downstream of Pax3 in NC induction and upstream of the other late NC-branch markers. This agreed with *sox9* being an early gene in the NB->NC branch. We tested the epistasis relationships between Pax3 and Sox9 in NC induction by combining Sox9 depletion or gain-of-function in the induced-neural crest assay (iNC is Pax3/Zic1-based NC induction from pluripotent ectoderm cells, Figure 6L). Early NC marker *Snail2* expression starts from gastrula stage 12.5 both *in vivo* and in iNC, and increases at neurula stage 14 (Figure 6 – figure supplement 1C). At both stages, co-activation of Sox9 strongly increased *snail2* expression in iNC, while Sox9 depletion reduced *snail2* activation (Figure 6L), indicating that Sox9 is required for efficient NC induction by Pax3 and Zic1. Interestingly, when iNC explants were analyzed prior to the normal onset of *snail2* expression, at mid-gastrula stage 11/11.5, Sox9 activation highly increased *snail2* expression, suggesting that Sox9 synergizes with Pax3 and Zic1 at the onset of NC induction.

In conclusion, we have defined a new transcriptional signature for the incompletely described Neural Border zone, established a global Ectoderm connectome and validated experimentally NC-related nodes, in particular highlighting how Sox9 enhances NC induction downstream of Pax3 and Zic1. We characterized three different transcriptional programs branching from neural, neural border and non-neural ectoderm progenitors towards NC and PC, and propose a model in which multiple co-existing paths lead to the early NC or placode states in gastrula-stage ectoderm.

## Discussion

In this work, we exploit the resolution of high-density single cell transcriptomes collected from 8 frog developmental stages to unravel the emergence of the neural crest lineage from the ectoderm during gastrulation, followed by diversification of neural crest progenitors during neurulation and upon EMT. Modeling gene transcription dynamics around each cell state allows the inference of the underlying molecular networks. We selected several important nodes for large-scale and *in vivo* experimental validation (Figure 7A, B). This study highlights the previously unknown temporal sequence of states in premigratory neural crest development. Firstly, we characterize neural crest activation either from a transient neural border state, or from a neural plate state during mid-gastrulation, and suggest a model that reconciles current debates upon multiple possible routes leading from immature ectoderm to neural crest and placodes (Figure 7C). Secondly, we delineate the early and later neural crest transcriptome trajectories during neurulation and define key regulators of branching, leading to eight transitional states and eight early migration states (Figure 7D).

**Figure 7.**
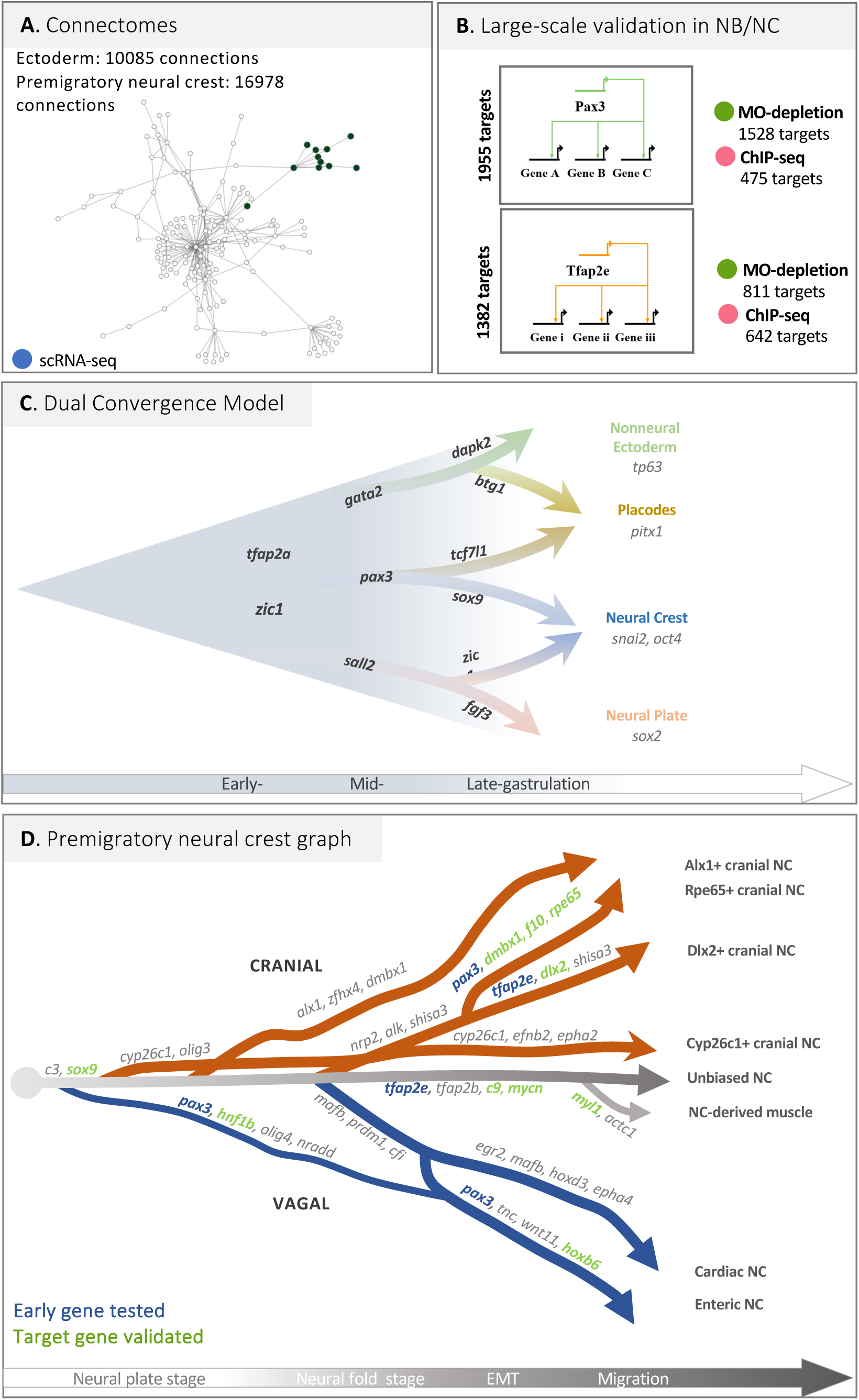
Neural crest GRN and developmental trancriptome trajectories. (A) GRNBoost2-predicted connectomes for Ectoderm (gastrula stage) and Neural crest (neurulation stage) can be queried at https://github.com/Qotov/neucrest_grn. (B) RNA-seq on dissected neural border ectoderm/neural crest explants and ChIP-seq provide large-scale GRNs linked to major nodes Pax3 and TFAP2e. (C) At gastrula stage (time line indicated), the neural border cells expressing tfap2a, zic1 and pax3 present trajectories towards placodes and neural crest. Those trajectories converge with a trajectory from neural plate towards neural crest and another one from non-neural ectoderm towards placodes. Branching analysis highlights a gene signature underlying those transcriptome transitions (the top gene is indicated here). (D) Transcriptomic tree of the neural crest cell states, from the end of gastrulation, during neurulation and upon epithelial-mesenchymal transition, ending at early migratory stage (time line indicated). Gene signatures supporting each trajectories are summarized. When Pax3 or Tfap2e were specifically expressed prior to branching (indicated in blue), their function on the expression of other genes in the branch-specific signature was tested in vivo. Genes with expression modulated after Pax3 or TFAP2e, manipulation in vivo and regulated are indicated in green.

### A reconciliatory “Model of Dual Convergence” describes the converging trajectories initiating neural crest and placode states

The molecular signature of the neural border ectoderm has been overtly simple, with only Pax3 (in frog and fish) or Pax7 (in chick) as relatively specific markers for this domain during gastrulation and early neurulation stages (Basch et al., 2006; Monsoro-Burq et al., 2005). Here, we have characterized the neural border ectoderm state by two features: the lower level of expression of genes expressed by adjacent neural (dorsal) and non-neural (ventral) cells (*sox2*, *lhx2*, *sall2*, *gata2, ker19*) and the increased expression of a large gene list including *zic1, tfap2a/c, pax3*, *sox9, hes1, cmyc*. We find that this state seems more transient than other ectoderm states, suggesting that these fate decisions occur quickly. In frog embryos, the end of gastrulation is clearly defined by blastopore closure, and this allows more precise exploration of timing compared to organisms with simultaneous gastrulation and neurulation such as chick embryos. In frog, fate choices in the dorsal ectoderm happen during the second half of gastrulation (between stage 11 and 12.5). As the neural plate forms and the blastopore closes (stage 13), fate decisions are clearly established between neural, non-neural, placode and neural crest with robust molecular signatures (Figure 6C). Modeling neural crest emergence from the neural border cell state confirmed the central role for Pax3 and suggested novel epistatic relationships between Pax3 and Sox9 upstream of the definitive neural crest state, defined by *snail2* expression. Importantly, we propose a novel model for the transcriptional pattern of decisions between the four main ectoderm fates, neural, non-neural, placodal, neural crest. Instead of contrasting “neural border” and “non-neural vs neural” hypotheses, we find that these routes are not exclusive and find trajectories supporting the emergence of neural crest from either the neural border or the nascent neural ectoderm on one hand, as well as two trajectories leading to placodes from either the neural border or the non-neural ectoderm. In each case, the gene programs underlying those alternative trajectories involve a subset of common genes and a few specific factors (Figure 6). For example, specific expression of *tcf7l1* and *stmn1* is found in placodes arising from NB zone, compared to the NNE route (Figure 6G, H). For neural crest, early *sox9* expression in the NB route contrasts with post-bifurcation expression in the NP route (Figure 6E, F). Thus, our SC transcriptome modeling reconciles and combines previously alternatives in a “Dual convergence Model” of neural crest and placode patterning, together with specific gene signatures ripe for future functional exploration (Figure 7C).

### A combination of Omics and in vivo strategies validates large sets of gene regulations driving the dynamics of neural crest diversification

The second outcome of our study is to define the temporal dynamics of trajectories that result into eight neural crest states present upon early migration stage along the cranial and vagal axial positions. The first key observation is the presence of a main population of NC unbiased towards any particular state, expressing markers of the immature neural crest cells, from which all the other trajectories emerge. This unbiased cell trajectory is maintained during and after EMT suggesting that a very plastic, stem-like NC cell population emigrates and is subjected to the signals from the microenvironment prior to fate choices. The second critical observation is that, for the anterior part of the body axis considered here, trajectories do not emerge in a spatially linear sequence from anterior to posterior as previously anticipated in a model where NC would follow an anterior-posterior wave of maturation. Two early trajectories arise from both the anterior (progenitors of posterior cranial NC - cluster 15) and the posterior-most positions (minor vagal trajectory progenitors, cluster 3) at neural plate stage (stages 13-14) prior to neural fold elevation. This is followed at mid-neurula stage (stage 15-16) by the emergence of the three other main cranial and vagal trajectories leading to cranial clusters 10 and 11, and to vagal NC clusters 12 and 13. Together with the maintenance of an immature stem-like cell population from induction to emigration, this sequence of trajectory determination suggests that the main cue controlling the temporal dynamics of states hierarchy in the cranial and vagal NC-GRN is not a function of the time elapsed since NC cell induction, or correlated to Hox gene positional information, but rather may involve response to external signals.

Our temporal analysis highlights three important points deepening our understanding of NC biology. Firstly, there have been long standing debates about the timing of NC fate decisions, prior or after EMT from the neural tube, in a variety of animal models (Kalcheim and Kumar, 2017). Importantly, we did not detect distinctive expression of predictive fate markers before EMT (e.g. for neuronal, glial skeletogenic or melanocyte fates). This suggested that, if some NC progenitors were biased towards a given fate prior to EMT, they did not exhibit a detectable signature in our dataset. However, our observations are in agreement with several lineage tracing studies showing the high multipotency of most NC cells when marked prior to EMT (Baggiolini et al., 2015). The first differentiation markers are found after emigration, as we detected myosin-like expression in a small subset of cells suggesting the emergence of previously poorly described NC-derived myofibroblasts shortly after EMT (Figure 2 – figure supplement 2). It also supports that our dataset is sensitive enough to detect other fate-specific markers if they were expressed at the end of neurulation. Secondly, our data support the early diversification into several distinct cell states prior, during and after EMT, contrasting with the recent suggestion that upon EMT the NC progenitors would regroup into a single common multipotent state (Zalc et al., 2021). The high cell content of our dataset proves otherwise, suggesting that this previous observation made on a smaller subset of cranial NC did not fully capture the diversity of pre-migratory NC states. Lastly, temporal trajectory analysis unravels the branch-specific dynamics of gene expression underlying bifurcations and state diversification. For each bifurcation, we provide a list of key genes likely to control branching choices (Figures 5 and 6). We further validate these predictions in several instances, by experimental modulation of pivotal transcription factors function in the premigratory neural crest (Pax3, TFAP2e), followed by *in vivo* or deep sequencing analysis. In sum, our study provides a comprehensive view of the hierarchy of molecular decisions driving the cranial and vagal neural crest gene regulatory network from induction at the neural border to early migration, with unprecedented resolution and deep learning-aided experimental validation. We propose a new “Dual Convergence Model” for neural crest and placode lineage emergence, and provide a detailed roadmap of the main molecular events in the premigratory and early migrating NC-GRN. Using a dedicated interactive network visualization interface, any gene of interest can be queried. Moreover, the detailed sequence of cell states provided here will prove an essential reference for monitoring induction of neural crest derivatives, for example from patient-derived induced pluripotent stem cells, when reliable specification protocols preferably recapitulate the steps of embryonic development.

## Materials ad Methods

### Experimental Design

Single cell transcriptomes from developing *X. tropicalis* embryos were scrutinized for NC development using machine-learning tools to infer the gene regulatory network (GRN) and the gene programs underlying branching of fates. These predictions were largely validated *in vivo* using micro-manipulations in *X. laevis* embryos followed by RNA-seq or ChIPseq. Detailed material and methods are given in Supplementary File 1.

### Single cell sequencing and processing

No new materials were collected for this study. Instead, we re-sequenced the SC RNA libraries for developmental stages NF11 to NF22 (Faber and Nieuwkoop, 2020) used in (Briggs et al., 2018) using NovaSeq S2. All datasets are deposited under NCBI Gene Expression Omnibus number GSE198494. For SC analysis, we used the *X.tropicalis* v.10 genome assembly, gene models v. 10.7, together with STAR aligner and the DropEst pipeline. After filtration by counts and genes numbers (>200 genes; >300 counts), we gathered a dataset of 177250 cells. In the cells of interest (Ectoderm and NC cells), mean counts number was 1778, and mean gene number was 1035. scRNA-seq postprocessing was done using Scanpy, a comprehensive scRNA pipeline that is functionally similar to Seurat (Stuart et al., 2019, see references and details in Supplementary File 1).

### Clustering and NC cells selection

For each stage, we obtained independent standard dimensionality reduction with PCA, computing a neighborhood graph and UMAP (Jacomy et al., 2014) followed by clustering, (Leiden algorithm, Traag et al., 2019). For each cluster, we defined cluster-specific genes with differential expression analysis (scanpy t-test_overestim_var) and selected only clusters which were the most similar to NC cells using NC signatures from (Briggs et al., 2018). For NC subclustering, we defined the optimal number of clusters by manually increasing their number and checking for biological meaning, as revealed by specific gene expression, for example of *hox* genes, such as *hoxd3* for cardiac NC (cluster #12), or *hoxb6* for Enteric Nervous System progenitors (ENSp, cluster #13).

### GRN generation

Using the temporal dynamics of gene expression, we used GRNBoost2 to infer genetic co-regulation, starting from a list of TFs (Blitz et al., 2017). GRNBoost2 retrieves the gene regulatory network (GRN) from the expression data-matrix (Moerman et al., 2019). In the resulting network of TFs and their targets we identified the most important nodes by calculating betweenness and degree centralities.

### Principal graph generation and branching analysis

The tree analysis was carried out using the scFates package (Albergante et al., 2020). However, this approach is too sensitive to build the principal graph for the whole NC dataset. Therefore, to generate the main tree, we used the PAGA algorithm (Wolf et al., 2018). This revealed cluster-cluster relationships including the early stages where the strongest connectivity was observed. Further we used ElPiGraph to study specific branches and bifurcation points. Using ScFates we defined features significantly changing along the tree, and then using pseudotime values and differential expression analysis, determined early and late branch-specific features. For each bifurcation point of interest, we selected a set of cells related to the clusters involved in the bifurcation. The selection of parameters for building the principal tree for each point of the bifurcation was carried out using brute force approach.

### Chromatin immunoprecipitation sequencing (ChIPseq)

Chromatin immunoprecipitation was performed according to (Wills et al., 2014) after injection of tracing amounts (75 pg) of mRNA encoding either Pax3-FLAG-HA or TFAP2e-FLAG). After sequencing, 100 bp single-end reads were aligned to *X. laevis* genome version 9.2 (Pax3) or *X. tropicalis* v10.0 (TFAP2e) using Bowtie2. Peaks were called using MACS2. For Pax3 we selected peaks common in three replicates, for TFAP2e we used stricter MACS2 score cutoff=500. Target genes were searched with bedtools (window size = 10kb).

### In vivo experiments: Xenopus laevis injections, microdissections, grafting and RNA-seq

*In vivo* injections, NB/NC dissections and grafting were done as previously (Milet and Monsoro-Burq, 2014; Plouhinec et al., 2017) using *X. laevis* embryos. For knockdown experiments, previously validated antisense morpholino oligonucleotides (MO) were used to deplete *pax3,* or *TFAP2e* transcripts (Monsoro-Burq et al., 2005; Hong et al., 2014). One Pax3 morphant anterior NB explant (stage 14) or one TFAP2e-morphant NC explant (stage 17) were dissected from the injected side, in triplicate. Each explant was sequenced individually (RNA-seq). The resulting 100 bp paired-end sequencing reads were aligned to the *X. laevis* genome version 9.2 using STAR and the count reads were analyzed using String Tie. Differentially expressed genes were selected considering log2FC and expression difference in absolute values (abs. diff. >=100 and <=500: log2FC>1.5 or log2FC<-1.5; abs. diff >=500 and <=1000: log2FC>1 or log2FC<-1; abs. diff >=1000 and <=3000: log2FC>0.5 or log2FC<-0.5; abs.diff.>3000: log2FC>0.33 or log2FC<-0.33)

### iNC assay

The induced neural crest assay (iNC) used co-activation of dexamethasone-inducible Pax3-GR and Zic1-GR at gastrula initiation stage 10.5, in pluripotent blastula ectoderm (animal caps dissected at blastula stage 9) (Hong and Saint-Jeannet, 2007; Milet et al., 2013). This was combined with Sox9 depletion (40 ng of *sox9* MO) (Spokony et al., 2002), or gain-of-function (300 pg *sox9* mRNA). At the desired stage, explants were harvested and processed for RTqPCR as in (Alkobtawi et al., 2021). Primers are listed in Table S8.

### Whole-mount in situ hybridization (ISH)

Whole-mount *in situ* hybridization followed a protocol optimized for NC (Monsoro-Burq, 2007). Embryos were imaged using a Lumar V12 Binocular microscope equipped with bright field and color cameras (Zeiss).

## Author contributions

Conceptualization: AK, SS, LP, AHMB

Methodology: AK, MA, SS, VK, SMR, HA, LP, AHMB

Investigation: AK, MA, SS, VK, SMR, HA, LP, AHMB

Visualization: AK, MA, SS, VK, AHMB

Supervision: LP, RMH, AHMB

Writing—original draft: AK, SS, AHMB

Writing—review & editing: AK, SS, RMH, AHMB

## Funding

This project received funding from

European Union’s Horizon 2020 research and innovation programme under Marie

Skłodowska-Curie grant agreement No 860635, NEUcrest ITN (AHMB)

Agence Nationale pour la Recherche (ANR-15-CE13-0012-01; AHMB)

Agence Nationale pour la Recherche ANR-21-CE13-0028; AHMB)

Institut Universitaire de France (AHMB)

National Institutes of Health NICHD award R01HD073104 (LP)

National Institutes of Health NIH R01 GM42341 (RMH)

National Institutes of Health NIH R35GM127069 (RMH)

## Acknowledgements

The authors are grateful to Drs. A. Zinoviev, G. Schlosser, I. Adameyko and T. Walter for insightful scientific discussions and comments on the manuscript. We thank C. Lantoine for animal husbandry, Q. Thuillier for technical assistance, and present and past members of the Monsoro-Burq lab for their support. We thank J. Briggs for help in single cell sequencing. We thank J.L. Plouhinec for preliminary RNAseq analysis. High-throughput sequencing, except for single cell sequencing, used the ICGex NGS platform of the Institut Curie supported by the grants ANR-10-EQPX-03 (Equipex) and ANR-10-INBS-09-08 (France Génomique Consortium) from the Agence Nationale de la Recherche (”Investissements d’Avenir” program), by the Canceropole Ile-de-France and by the SiRIC-Curie program -SiRIC Grant « INCa-DGOS-4654 ».

## Data and materials availability

All data are available in the main text or the supplementary materials. Biological reagents are available upon request to the corresponding author. Accession numbers to the datasets, together with their description are under NCBI Gene Expression Omnibus # GSE198494.

## List of Supplementary Materials

- Figure 1 – figure supplement 1
- Figure 2 – figure supplements 1, 2, 3, 4 and 5
- Figure 4 – figure supplements 1 and 2
- Figure 5 – figure supplement 1
- Figure 6 – figure supplement 1
- Supplementary File 1

a. Tables S1, S2 and S3
b. Detailed description of NC clusters
c. Tables S4, S5, S6, S7 and S8
d. Supplementary Matdrials and Methods

---

**Figure 1 - figure supplement 1.**
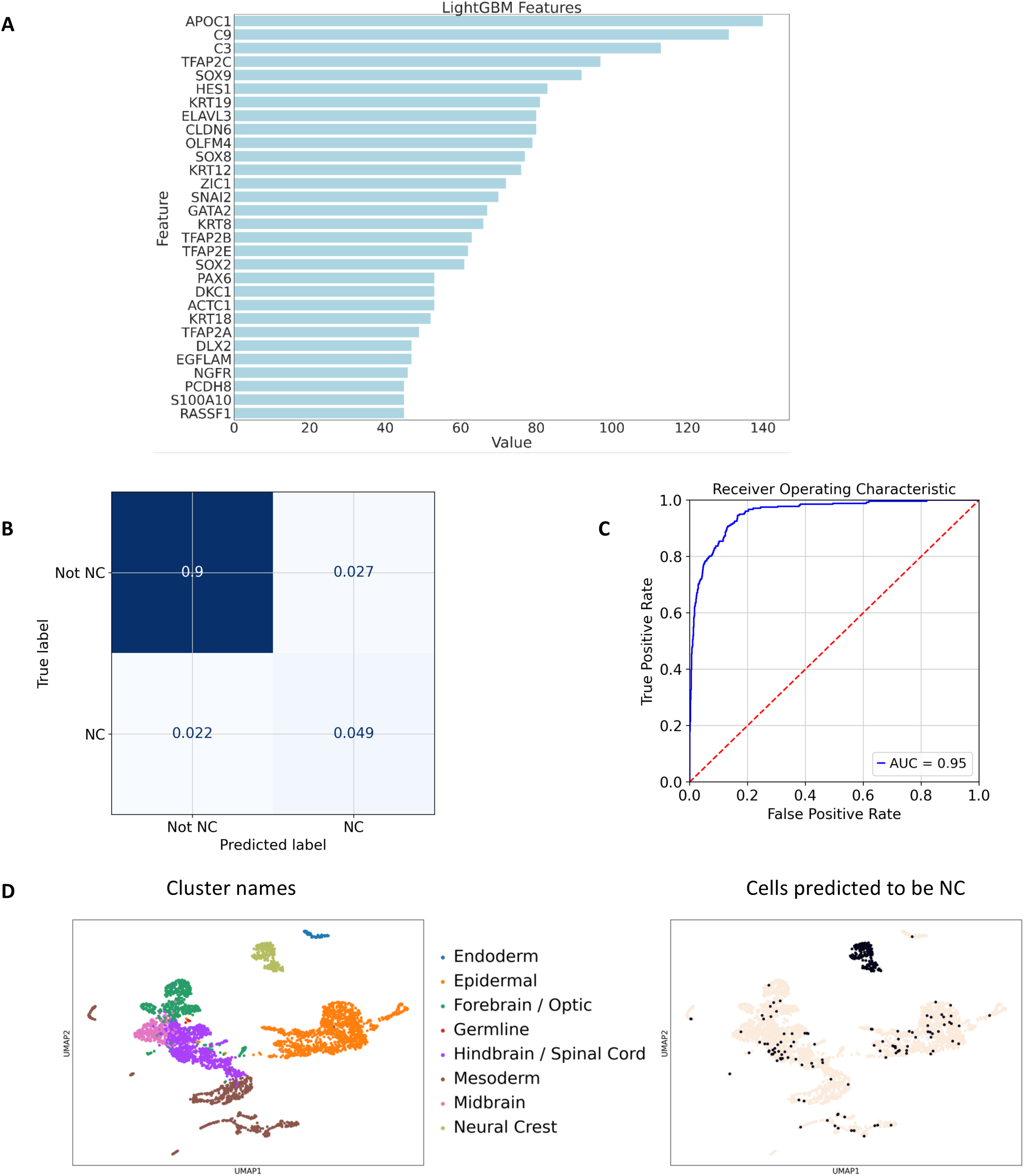
A NC classifier based on Xenopus NC selection retrieves NC cells in other whole embryo datasets. A binary classifier was trained on the frog NC/non-NC cells (see methods). Using the whole embryo data matrix of gene expression, we selected the top 10% important features for NC detection. In order to validate it in a whole embryo SC Zebrafish dataset (Wagner et al., 2018), we removed genes present in frog data that were absent in the Zebrafish dataset: Using the subselected top genes, we re-built the model using frog data and obtained the following results: 0.99 accuracy and 0.90 f1 score on a test dataset. When applied to Zebrafish dataset, we obtained also very high scores: the accuracy 0.95 and f1 score 0.66 for 14 stage cells, indicating good initial NC cell selection. (A) Top features (gene) for NC classification using LightGBM model. X axis value is the relative importance of the certain gene for the NC classifier (B) Confusion matrix indicates the true/false positive and the true/false negative scores for the results obtained on the Zebrafish dataset for stage 14. (C) ROC curve for the binary NC classifier indicates high AUC score of 0.95, meaning high recovery of true positives by the model. (D) UMAP for Zebrafish stage 14 hpf cells (left) with model-predicted NC cells. The model accurately identifies the cluster of interest (NC-khaki) despite the technical and biological batch effects between Xenopus and Zebrafish datasets.

**Figure 2 - figure supplement 1.**
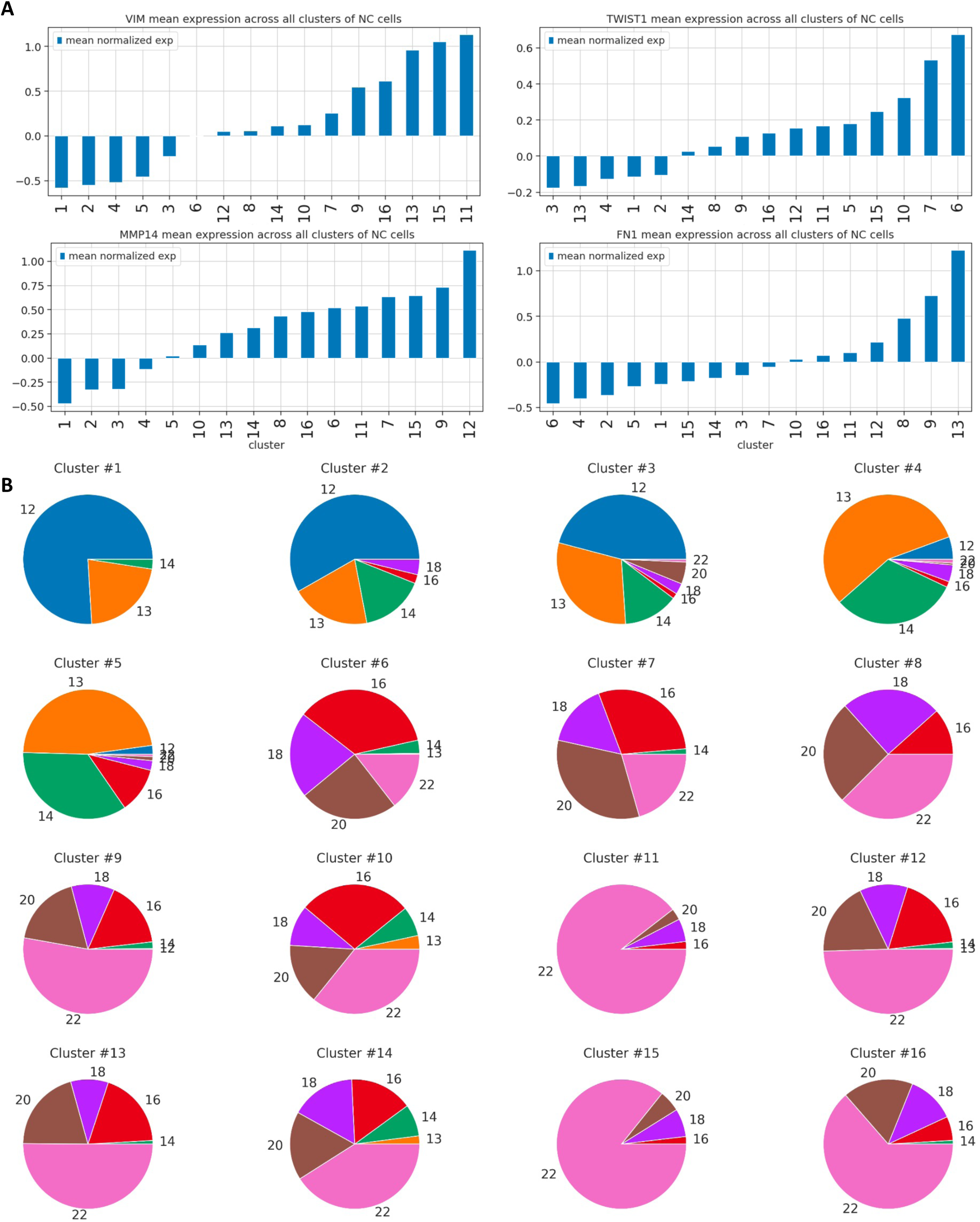
Developmental stages and expression of EMT/migration markers per cluster. (A) Each cluster presents various levels of expression of EMT and migration regulators. Bar plots with mean normalized expression for each cluster for vim, twist1, mmp14 and fn1. (B) Pieplots represent the proportion of cells at each stage in each cluster. Some clusters include cells of 6 successive stages, e.g. cluster 10.

**Figure 2 – figure supplement 2.**
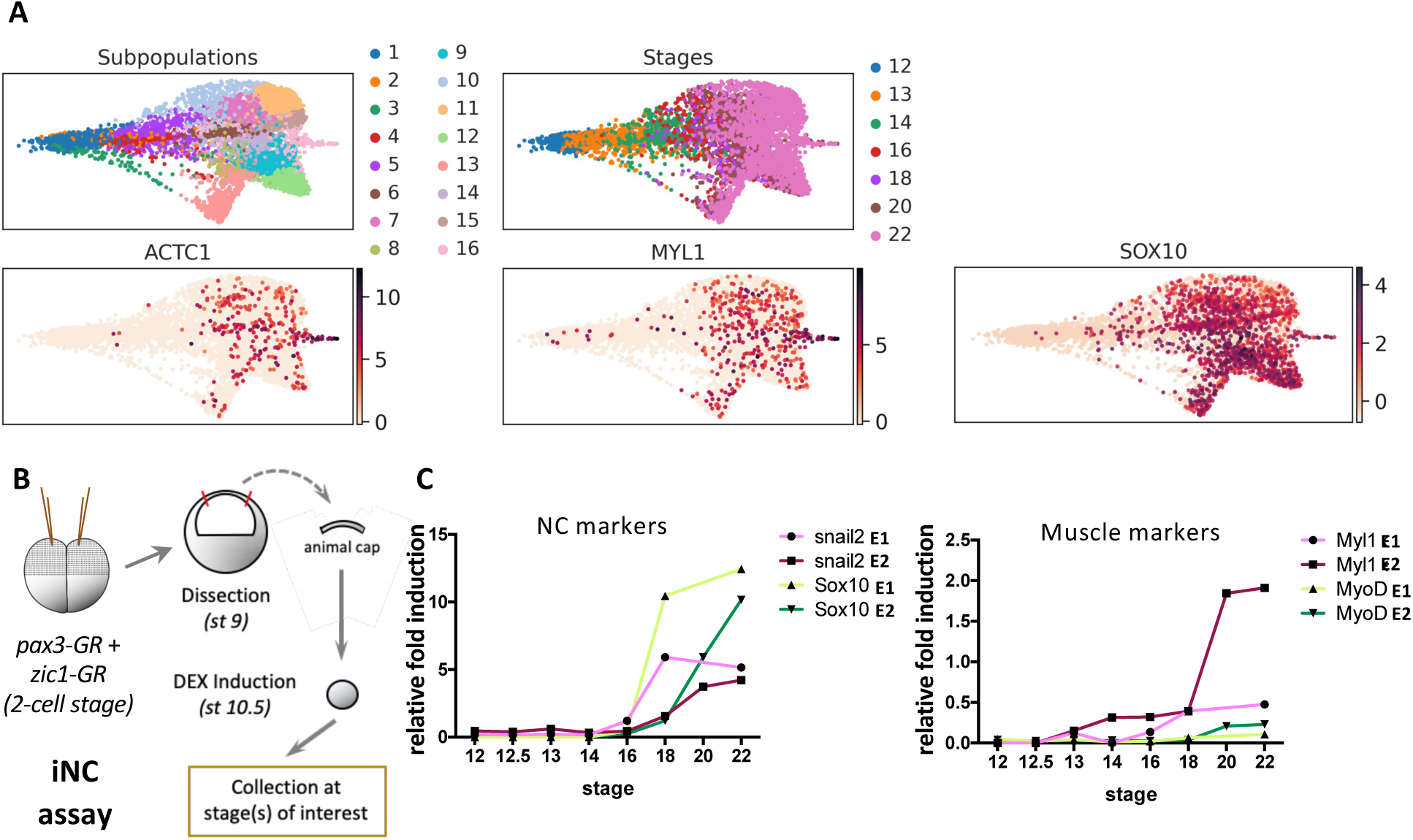
Identification of a muscle-like cluster. (A) At early migration stage (20-22), cluster 16 is strongly enriched for myosin-like genes as well as bona fide NC markers, such as sox10. (B, C) Direct NC induction in ectoderm explants grown in vitro (iNC assay) followed by RTqPCR from induction to emigration stages, also identifies the late activation of myoD and myl1 expression at stage 20. This validates the appearence of this muscle-like NC subpopulation among other NC derivatives.

**Figure 2 - figure supplement 3.**
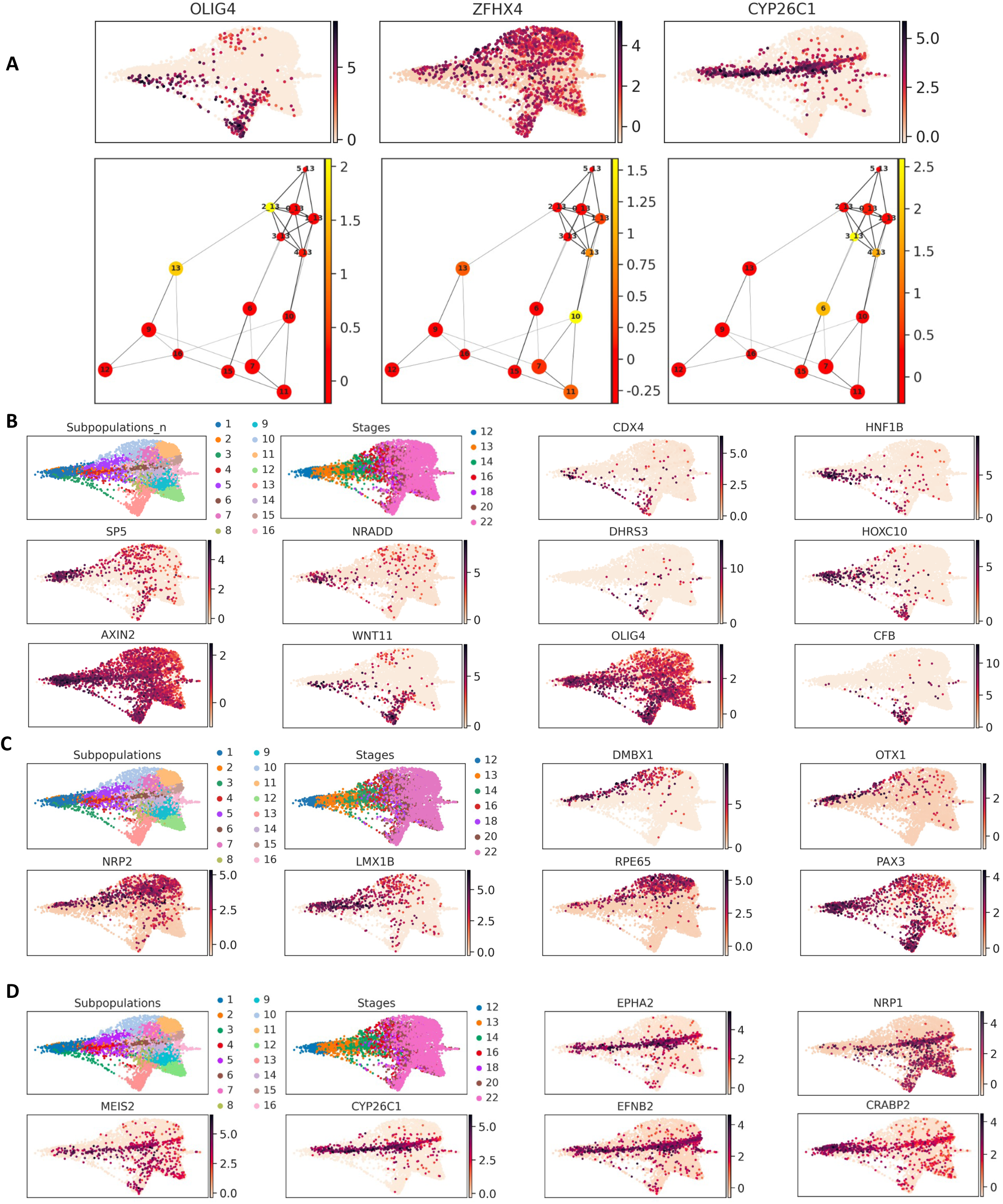
Early stage biased NC clusters. (A) From the end of gastrulation, three previously undescribed early trends emerge. Clustering of stage 13 cells only, followed by PAGA analysis identified their similarities with the stage 20-22 clusters: an olig4-enriched subpopulation linked to ENSp cluster 13; a zfhx4-enriched cluster related to rpe65+ cluster 10 and a third stage 13 subpopulation expressing cyp26c1 linked to the cranial NC branch (clusters 4, 6, 15). (B) UMAP plots for the early-biased NC subpopulation expressing hnf1b, cdx4, nradd and posterior hox genes. In addition to cluster-specific hnf1b and olig4, dhrs3 is co-expressed with zic5 which specifically stays active during the whole “sacral” trajectory from stage 12 to late ENSp. (C) UMAP plots of genes specific for cranial rpe65+hox- NC subpopulation specific genes (D) UMAP plots of genes specific for rhombomere 4, cranial NC subpopulations.

**Figure 2 - figure supplement 4.**
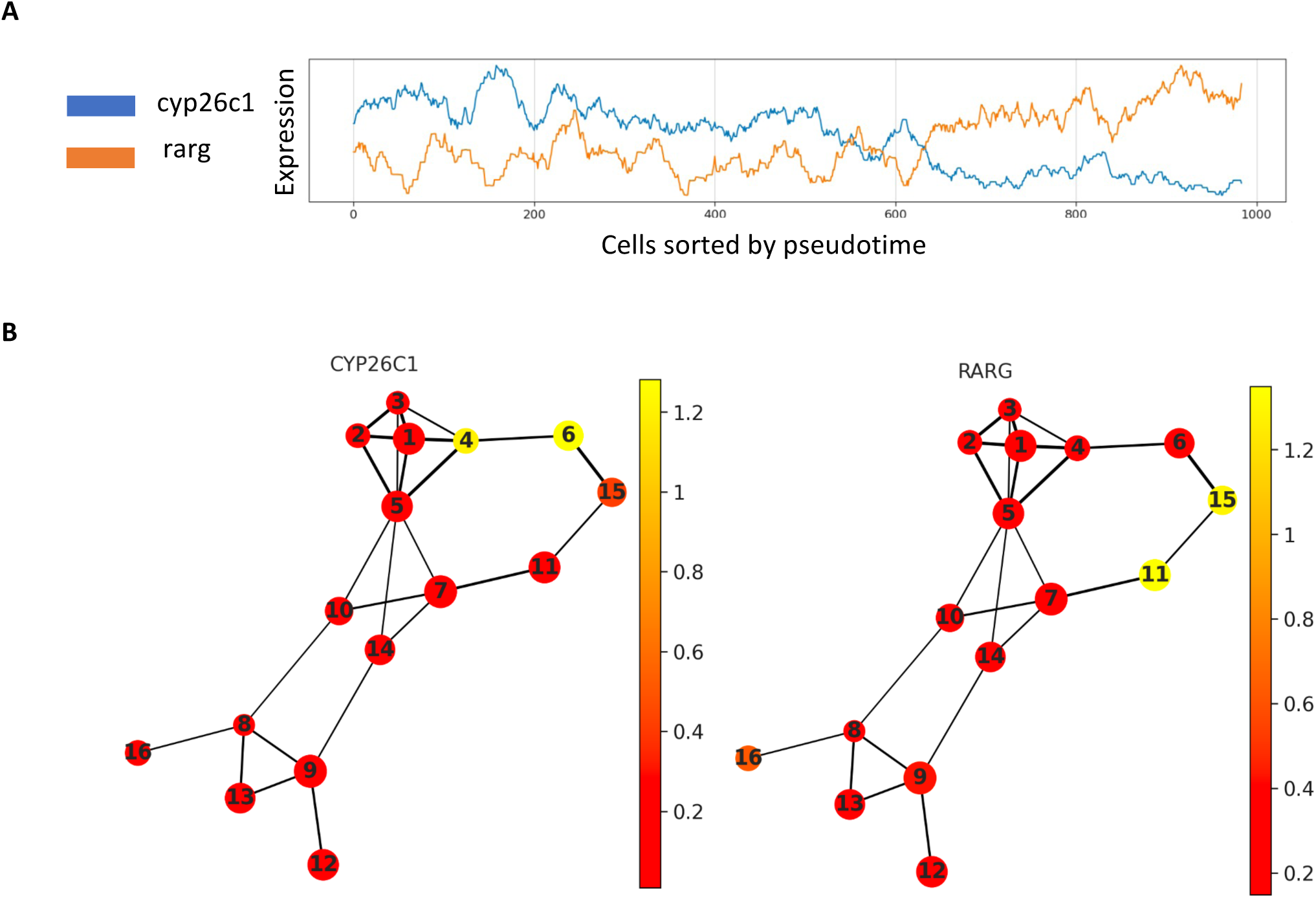
Early posterior enteric cluster 4 and related late clusters 6, 15. (A) Line Plot for expression dynamics for cluster 4, 6, 15, representing the relationships between rarg and cyp26c1. Cells within clusters are sorted according to pseudotime. (B) PAGA plots with rarg and cyp26c1 mean expression.

**Figure 2 - figure supplement 5.**
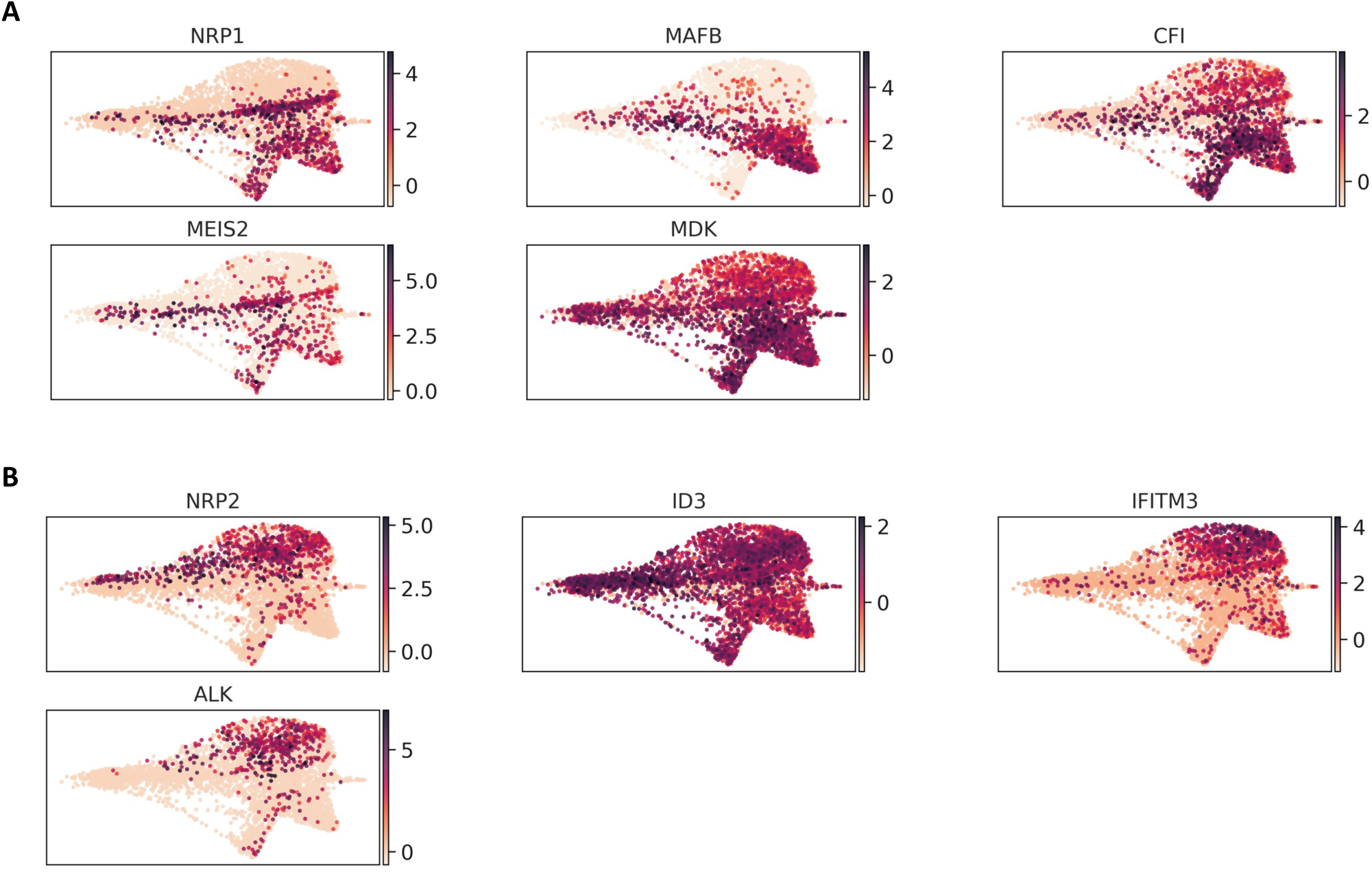
Early stage, bipotent NC clusters – cranial vs vagal lineages. UMAP plots for vagal (clusters 9, 12 and 13) and cranial (clusters 7, 10 and 11) specifiers.

**Figure 4 - figure supplement 1.**
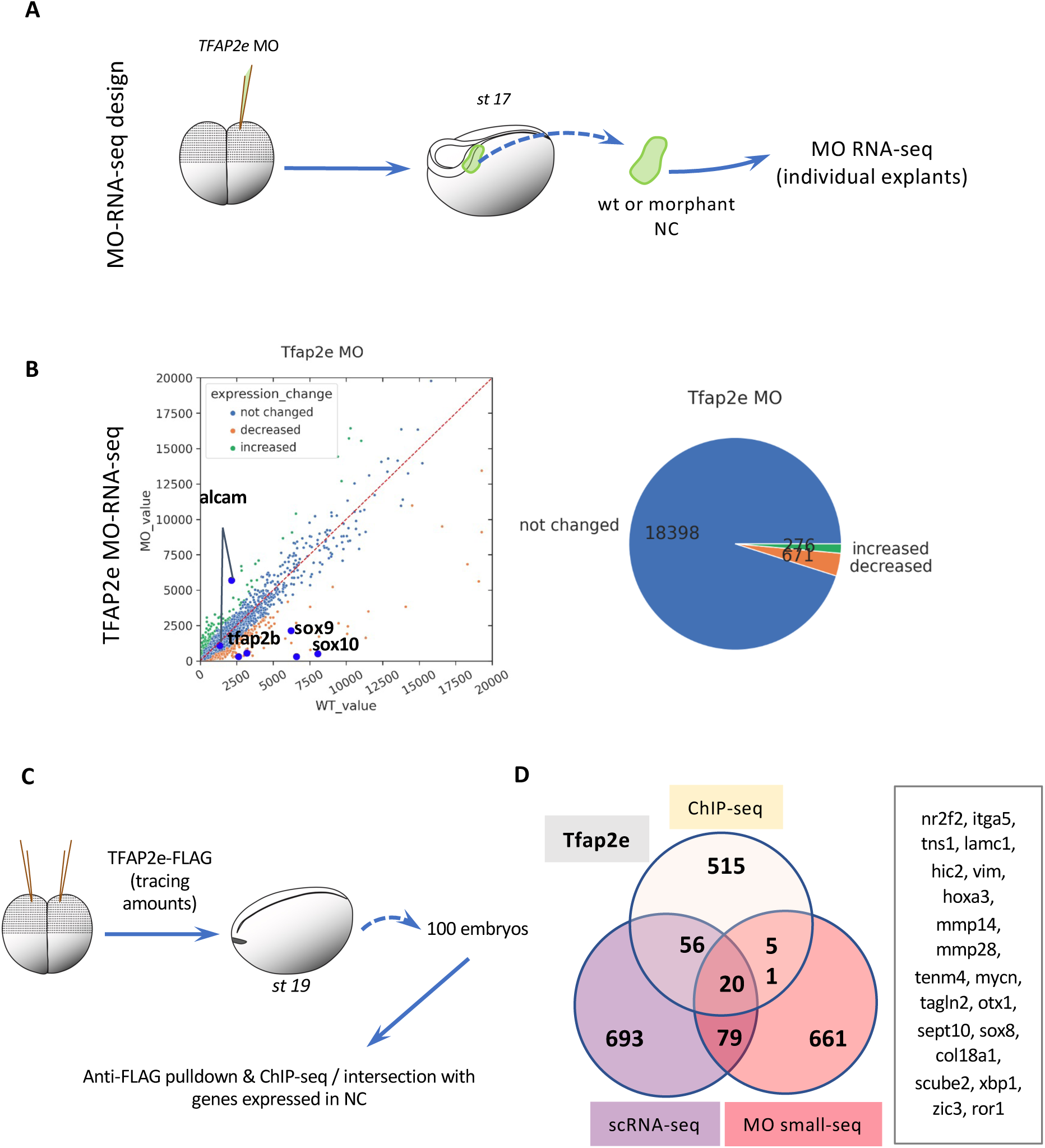
TFAP2e connectome generation and validation. **(A)** Design of RNA-seq on microdissected NC explants after depletion of TFAP2e. **(B)** General statistics for differential analysis after RNA-seq of NC depleted for TFAP2e. **(C)** Design of ChIP-seq experiments for TFAP2e. **(D)** Venn diagram of target genes validated by ChIP-seq and transcriptome of TFAP2e-depleted NC, compared to predictions using GRNBoost2. The 20 genes linked to TFAP2e by all three methods are listed, which represents a significant increased compared to a “at random” situation. Among them we find genes important for NC EMT and migration *mmp14, mmp28, itga5* and *vim*.

**Figure 4 - figure supplement 2.**
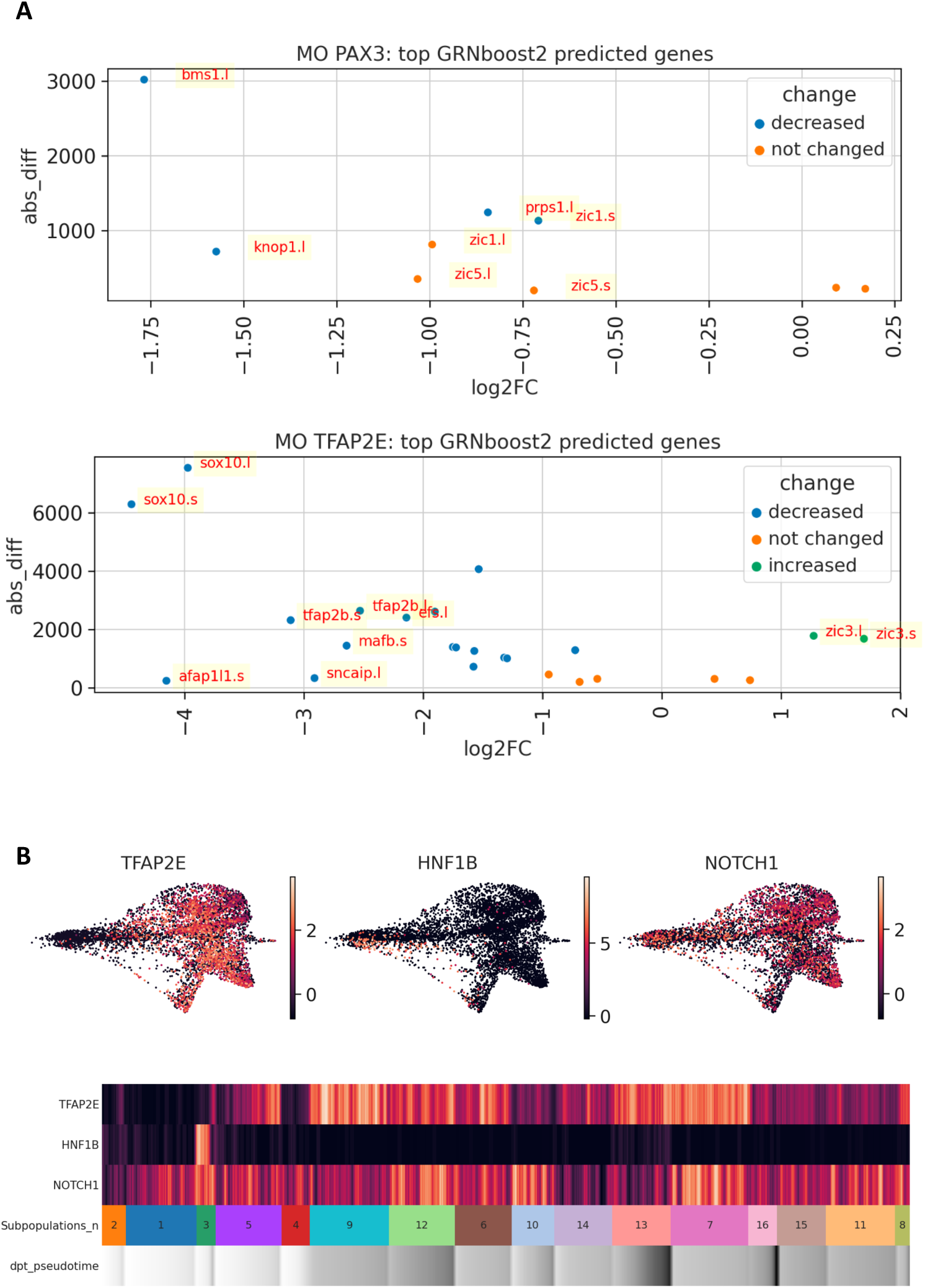
Validation of Pax3 and TFAP2e connectomes. (A) The expression levels of the genes top-scored with GRNboost2 for Pax3 or TFAP2e were tested after Pax3 or TFAP2e depletion in NB/NC *in vivo*. Several of them were strongly decreased after Pax3 or TFAP2e depletion (e.g. *bms1* or *sox10* respectively). Others were unchanged meaning that, despite similar transcriptome dynamics at the single cell level (evaluated by GRNBoost2), there was no major functional regulation by either Pax3 or TFAP2e *in vivo* at the considered stage (unless unknown compensatory mechanisms were to apply). (B) Conversely, we identified some targets of TFAP2e, confirmed with ChIP-seq and MO, which were not predicted by GRNboost2 (e.g. *hnf1b* and *notch1*). This may result from functional regulations creating different expression patterns or different temporal dynamics, or else expression in adjacent cells (non cell-autonomous regulations), which are all parameters for which GRNBoost2 would not identify linkage. For example, neither *hnf1b* nor *notch1* were predicted as TFAP2e targets: *hnf1b* displays a very different expression pattern than *TFAP2e,* in the NC dataset, while *notch1,* despite a closely-related expression pattern at the cluster level, may in fact display distinct cell-to-cell expression at the single cell level.

**Figure 5 - figure supplement 1.**
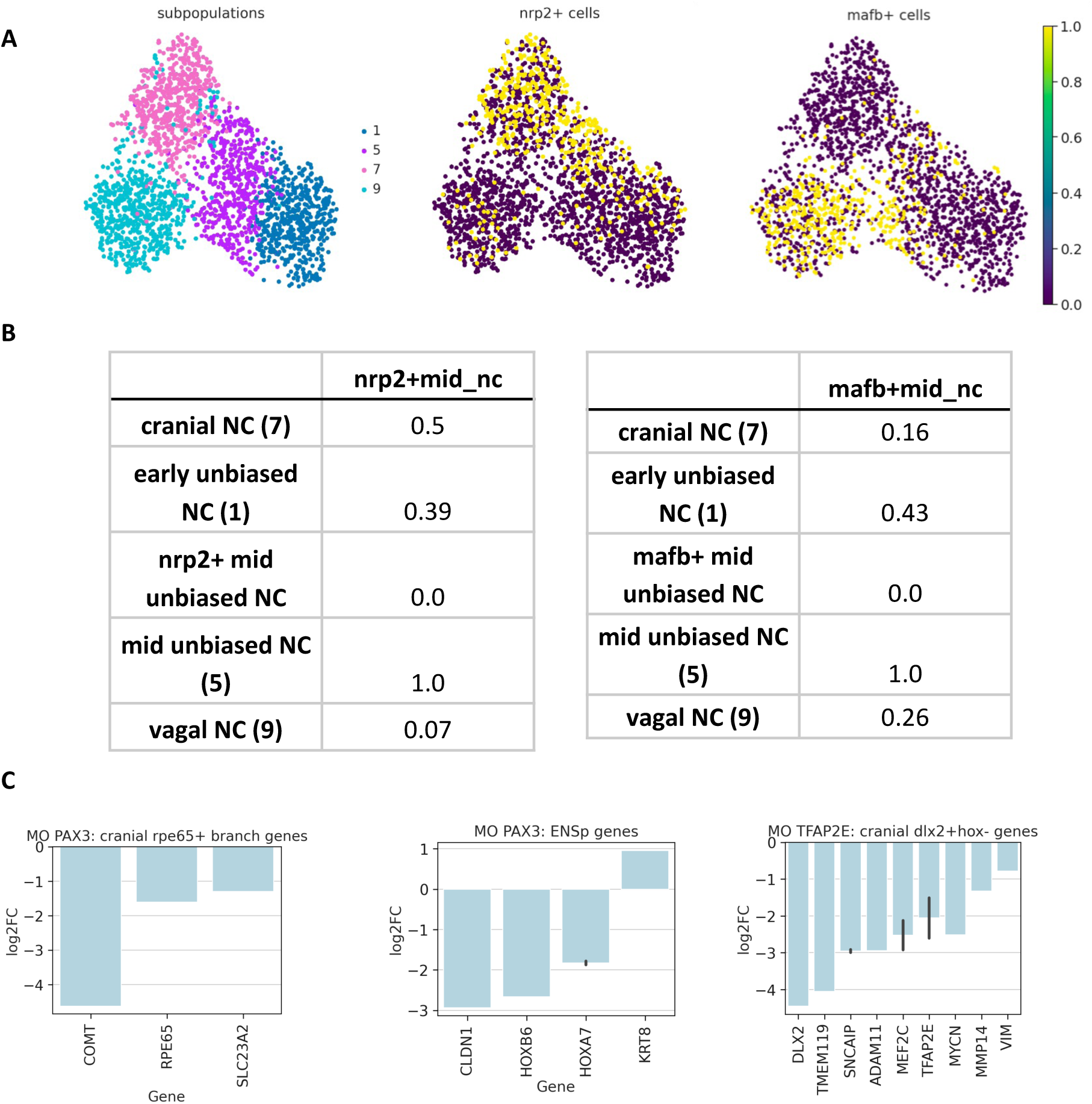
Branching analysis for NC dataset. **(A)** In order to identify potential early bias in the globally homogeneous cluster 5, cells located around the bifurcation between clusters 1 (dark blue), 5 (purple), 7 (pink), and 9 (cyan) were sub-selected. Cells expressing *nrp2* or *mafb* above 80 percentile were highlighted. This revealed that cluster 5 presented a minor predisposition for either *mafb* or *nrp2* expression. **(B)** Tables depicting cluster-cluster similarities calculated with PAGA. PAGA revealed that *nrp2*^+^ cells of cluster 5 were 7 times more similar to the cranial state (cluster 7) than to the vagal state (cluster 9). Conversely *mafb*^+^ cells of unbiased cluster 5 were 1.5 times closer to the vagal state than to the cranial. **(C)** To test branching analysis predictions, we used *in vivo* depletion of either Pax3 or TFAP2e, and tested the changes in expression of the late genes in Pax3- or TFAP2e-related branch.

**Figure 6 - figure supplement 1.**
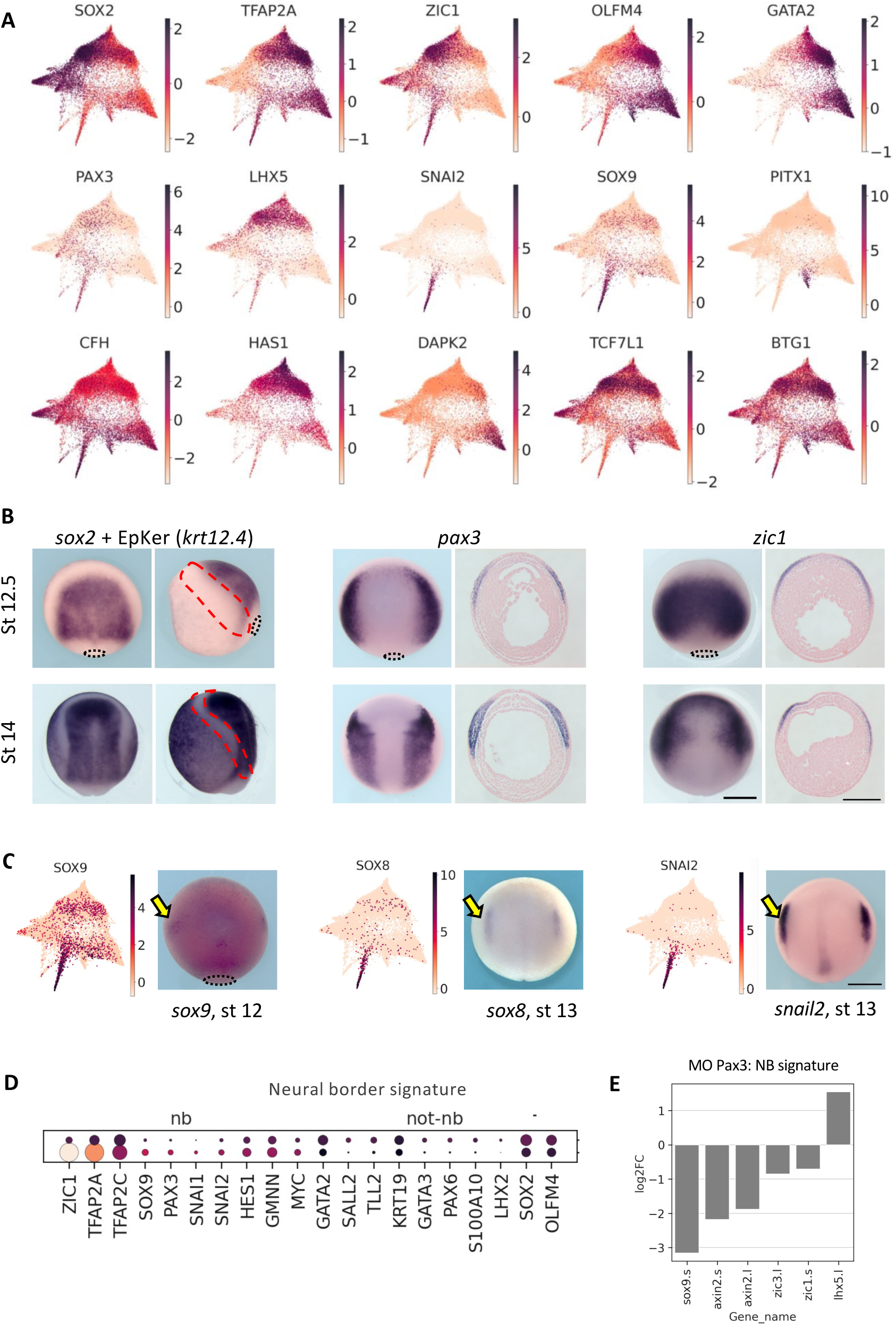
Markers for NB zone and early NC. **(A)** Ectoderm UMAP plots for stages 11, 12 and 13 representing gene expression of the main markers for ventral, dorsal and neural border, across the gastrula stage ectoderm. **(B)** In situ hybridization patterns for genes marking the 3 major ectoderm areas at stages 12.5 (late gastrula) and 14 (neurula). On dorsal and lateral views, *sox2* marks the neural (dorsal) ectoderm while *krt12* (epidermal keratin 12) marks the non-neural (ventral) ectoderm. Unlike the other derivatives, the NB zone does not yet have a large and specific gene marker list and is usually depicted by *pax3* expression or the overlapping expressions of dorsal and ventral genes like *zic1* and *tfap2a* respectively (dorsal views and transverse histological sections after whole mount *in situ* hybridization). Importantly, the neural border area (red dotted line) can also be defined as a zone devoid of both *sox2* and *keratin* expression as early as stage 12.5. **(C)** Ectoderm UMAP and corresponding ISH images depict the earliest stages of detection of 3 well-known early NC markers. *Sox9* is initiated at the prospective NC zone at stage 12. *Sox8* and *snail2* are detected at the early NC at stage 13. Yellow arrows point to the NC region. Scale bar, 500 *¼*m. **(D)** Heatmap identify new genes enriched in the neural border defining an enlarged gene signature. **(C)** Neural border genes expression affected after Pax3 depletion *in vivo*.

