## Supplementary materials for "From neural border to migratory stage: A comprehensive single cell roadmap of the timing and regulatory logic driving cranial and vagal neural crest emergence"

**Supplementary File 1**

**From neural border to migratory stage: A comprehensive single cell roadmap of the timing and regulatory logic driving cranial and vagal neural crest emergence**

Aleksandr Kotov, Mansour Alkobtawi†, Subham Seal†, Vincent Kappès, Sofia Medina, Hugo Arbès, Richard Harland, Leonid Peshkin, and Anne H. Monsoro-Burq^*^

**Contents**

- Table S1: Comparison of this Ectoderm/NC dataset with the relevant previously published scRNAseq datasets related to early NC development.
- Table S2: Top Ectoderm and NC specific genes for each developmental stage.
- Table S3 and associated text: Premigratory NC clusters characterisation and key enriched genes.
- Supplementary Figure Legends
- Table S4 & S5. Top-decreased genes after Pax3 or TFAP2e2e depletion.
- Table S6. Pax3 targets genes identified by ChIP-seq.
- Table S7. TFAP2e ChIP-seq targets.
- Table S8. PCR Primers used in this study.
- Supplementary Materials and Methods
- Supplementary references

**Table S1.**

**Comparison of this Ectoderm/NC dataset with the relevant previously published scRNAseq datasets related to early NC development.**

Compared to previously published SC transcriptome datasets of NC cells, our new dataset includes the largest number of premigratory and early-migrating NC cells. This allowed us to use complex ML approaches to reveal gene regulatory relationships.

| Dataset | NC selection | Cell number | Depth | Timing | scRNA-seq technology | Reference |
| --- | --- | --- | --- | --- | --- | --- |
| GSE113074 | X.tropicalis WE/markers | 130000+ WE, 4700 NC | Low | 9 gastrula and neurula batches | Indrops v2/v3 | Briggs et al, 2019 |
| GSE112294 | Zebrafish WE/markers | 63520 WE, 1770 NC | Low | 7 gastrula & neurula batches | Indrops v1 | Wagner et al, 2018 |
| GSE106676 | Zebrafish, foxd3+ cells | 96 NC | High | 1 gastrula & 3 neurula stages | SmartSeq2 | Lukoseviciute et al, 2018 |
| GSE130500  GSE131688 | Chick, foxd3+ | 137 SmarSeq2  3000 10x | High + Low | Late neurula | SmartSeq2  10x | Williams et al, 2019 |
| GSE129114 | Mouse, wnt1/sox10+ | 200 NC | High | Neurula | SmartSeq2 | Soldatov et al, 2019 |
| GSE162044 | Mouse, wnt1+ NC | 1741 NC | High | Neurula | Smart-Seq2 | Zalc et al, 2021 |
| GSE198494 | X.tropicalis WE/markers | 140000+ WE, 6135 NC  17138 EC | Low | 7 gastrula and neurula batches | Indrop v3 | This study |

**Table S2. Top Ectoderm and NC specific genes for each developmental stage.**

Using Leiden clustering and differential expression analysis we constructed two datasets: Ectoderm and NC. Table represents the most specific genes expressed in the clusters in comparison to other cells of the whole embryo.

| Cell type | NF stage | Cells number | Gene markers for cluster characterisation |
| --- | --- | --- | --- |
| EC | 11 | 9190 | SOX2, OLFM4. FOXI1, GATA2, SOX11 |
| EC | 12 | 3500 | TFAP2C, C3, GATA2, OLFM4, ZEB2 |
| EC | 13 | 4448 | TFAP2C, SOX9, OLFM4, LHX2 |
| NC | 12 | 623 | C3, SOX9, ZIC1, SNAI2 |
| NC | 13 | 579 | SOX9, SNAI2, C3, ZIC1 |
| NC | 14 | 416 | SNAI2, TFAP2B, SOX9, SOX8 |
| NC | 16 | 882 | TFAP2B, C9, C3, SOX10 |
| NC | 18 | 607 | TFAP2B, C9, SOX8 |
| NC | 20 | 839 | TFAP2B, SOX8, C9 |
| NC | 22 | 2253 | TFAP2B, C3, C9, SOX10, DLX2 |

**Premigratory NC clusters characterisation and key enriched genes.**

**Table S3.**

| **Cluster number** | **Characteristics** | **Specific marker gene** |
| --- | --- | --- |
| 1 | Early unbiased | *tfap2c^+^* |
| 2 | Early unbiased | *gjb1^+^* |
| 3 | Premigratory sacral | *hnf1b^+^* |
| 4 | Premigratory early R4 | *cyp26c1^+^* |
| 5 | Premigratory unbiased | *-* |
| 6 | Premigratory late R4 | *cyp26c1^+^* |
| 7 | Migratory bipotent cranial | *nrp2^+^* |
| 8 | Migratory rhombencephalic | *hoxb2^+^* |
| 9 | Migratory bipotent vagal | *hoxd3^+^* |
| 10 | Premigratory early cranial | *alx1^+^* |
| 11 | Migratory cranial | *itga4^+^* |
| 12 | EMT cardiac | *egr2^+^* |
| 13 | Migratory ENSp | *tnc^+^* |
| 14 | EMT unbiased | *-* |
| 15 | Migratory late R4 | *efnb2^+^* |
| 16 | Migratory muscle-like | *myl1^+^* |

**Detailed description of the premigratory and early migrating neural crest clusters**

Out of ten migratory stage clusters (7-16), we identified five cranial NC cell subpopulations based on the lack of *hox* gene expression and expression of known anterior NC markers. Of these, three were previously found in (Briggs et al., 2018): *rpe65^+^* NC cluster 10; *vim^+^, itga4^+^* enriched NC cluster 11; and *dlx2^+^, efnb2^+^, cyp26c1^+^, epha2^+^* NC cluster 15. We uncovered two previously undescribed clusters: cluster 16, a muscle-like NC subpopulation, enriched for actin and myosin-like genes (*actc1^+^, myl1^+^* cells) (Figure 2 – figure supplement 2A), and cluster 14, a population of NC cells expressing the canonical early neural crest signature (*tfap2b, c9*) without additional specific signature, suggesting that these cells, although adopting mesenchymal characteristics (*vim* expression), remain unbiased and potentially multipotent (Figure 3).

We used the well-established induced neural crest (iNC) assay in *Xenopus laevis* embryos to validate the NC origins of cluster 16 (Figure 2 – figure supplement 2B, C). In this assay, NC cells are induced from pluripotent ectoderm, devoid of mesoderm contribution (confirmed by the absence of early myoD expression). We found that *myl1* and *myoD* started being expressed around stage 20 from the induced NC progenitors, indicating that this muscle-like NC cell population is also induced among the other derivatives described for the iNC differentiation assay (Milet and Monsoro-Burq, 2012). This population may correspond to early NC-derived head muscles or myofibroblasts (Grenier et al., 2009).

Additionally, we identified three subpopulations of *hox-*positive rhombencephalic neural crest previously grouped together (Briggs et al., 2018). These cells form clusters 12, 13 and 9 and express *hoxa3, hoxb3, hoxd3* respectively*,* and are thus located at the vagal level. Cluster 12 includes cardiac NC (*egr2^+^, mafb^+^*) (Tani-Matsuhana and Inoue, 2021) while cluster 13 contains (*tnc^+^, ltbp1^+^*) ENS progenitors (Akbareian et al., 2013) (Figure 2D, E).

Using specific gene signatures for clusters 1-6, we described the main characteristics of progenitors for each later stage NC subpopulation (clusters 7-16). From the earliest stage of NC induction (gastrula stages 12-13), we detected three distinct trajectories emerging from the main canonical NC progenitor population (clusters 1, 2): an *hnf1b*^+^*olig4*^+^ cluster 3, a *cyp26c1*^+^ cluster 4 and a *rpe65*^+^*zfhx4*^+^ cluster 10 (Figure 2E, Figure 2 – figure supplement 3A). First, the previously undescribed *hnf1b*^+^*olig4*^+^ cluster 3 is a pre-migratory cluster composed mainly of stage 12, 13 and 14 cells expressing pan-NB/NC markers such as *pax3, tfap2b* and *sox9,* as well as the dorsal neural gene *olig4, hnf1b,* the *ngfr-*related gene *nradd*, complement factor *cfb,* the posterior master regulator *cdx4* and a strong posterior hox gene signature (including posterior hox gene *hoxb8* and sacral hox genes *hoxc10/hoxd10*) (Figure 2D, Figure 2 – figure supplement 3B)*.* Interestingly, this cluster seemed to respond to a combination of high WNT/low RA signals, in line with its posterior trunk signature (Martínez-Morales et al., 2011). Indeed early on, cluster 3 expressed canonical WNT signaling targets *axin2* and *cdx4* (Kjolby and Harland, 2017), as well as *dhrs3*, which attenuates RA signaling, is required for posterior axis formation and is usually mutually exclusive with *cyp26c1* expression (Kam et al., 2013). In cluster 3’s older cells, expression of genes responsive to canonical WNT signals, *cdx4* and *sp5,* was decreased, while expression of the non-canonical ligand *wnt11* was increased. Cluster 3’s trajectory converged and merged with the enteric NC cells of cluster 13. While most of the enteric peripheral nervous system (ENS) arises from vagal NC, sacral NC cells contribute to cells in the hindgut. Cluster 3 posterior gene signature suggested that as early as mid-gastrulation stage 12, a minor NC progenitor population is already biased towards the future sacral NC fate, although those cells would emerge later on from the neural tube at tailbud-stage and give rise to the posterior ENS (Burns et al., 2004; Collazo et al., 1993; Sasselli et al., 2012). Moreover, *wnt11* negatively regulates enteric NC differentiation (Nagy et al., 2021), further suggesting that cluster 3 trajectory may consist of immature ENS progenitors. This cluster is the most posterior cell group found in this dataset (posterior trunk/sacral *hox* signature). Identification of cluster 3 specific markers will enable future lineage tracing experiments to define the position and developmental contribution of these early progenitors of the enteric nervous system.

Appearing at stage 13, the early-biased *hox*^-^ cluster 10 expresses *dmbx1, otx1, nrp2, lmx1b, rpe65* and *pax3,* thus contains early premigratory cranial NC cells (Figure 2 – figure supplement 3C). Bias towards cluster 10 was slightly noticeable at stage 12, as around 5% of the *unbiased* cluster 1 co-expressed *dmbx1* and *otx1*, which might indicate early fate predetermination. Moreover, 10% of the cluster 10 were cells of 13-14 stages (Figure 2 – figure supplement 1B), the rest being later stages. This means that progenitors for cranial NC start being specified at neural plate stage, much earlier than anticipated.

The third early-biased cluster, cluster 4, was composed of neural plate stage 13-14 progenitors of cranial NC populations with a unique retinoic acid (RA) signaling-related signature. It merged with cluster 6, (composed of stage 16-20 cells) and cluster 15 (composed of stage 22 cells). Cluster 4 specifically expressed genes encoding the RA-degrading enzyme *cyp26c1,* transcription factors *meis2* and *olig3,* neuropilin receptor *nrp1,* ephrin signaling receptor *epha2,* and ligand *efnb2* (Figure 2 – figure supplement 3D)*.* Compared to the early unbiased cluster 1, cluster 4 exhibited increased expression of mature pan-NC markers *tfap2b, c9, snai2*, and decreased expression of multipotency marker *pou5f1* (Figure 3B). Cluster 4 expressed *hoxb1* and *hoxb2* indicating an anterior rhombencephalon position. More precisely *epha2* expression pinned those cells to rhombomere 4 (Figures 2D) (Helbling et al., 1998). High *cyp26c1* expression indicated low RA levels as essential to define rhombomere 4 identity (Addison et al., 2018)**.** Last, *nrp1* specifically marks mouse NC from rhombomere 4 suggesting high conservation of the cluster signatures across vertebrates (Lumb et al., 2014). Cluster 4’s *cyp26c1/epha2* signature remained specifically expressed in the low-RA-dependent stream (6, 15) from stage 12 to 22. Expression levels of genes encoding *epha2* and *efnb2* remained high while *cyp26c1* expression gradually decreased in later subpopulation of posterior CNC along with increased expression of genes encoding RA receptors (e.g. *rarg*) and cellular retinoic acid-binding protein 2 (*crabp2*) (Figure 2 – figure supplements 3D and 4A, B)*.* This revealed that the identity of rhombomere 4 NC was established as early as neural plate stages 13-14 under low RA signaling (cluster 4) while starting at neural fold stage 16 (cluster 6), later development of this lineage involved increased RA signaling. These observations highlighted the transcriptional outcomes of dynamic RA signaling in the progenitors of rhombencephalic neural crest forming the first (mandibular) and second (hyoid) arch stream respectively, anterior to the otic vesicle.

Interestingly at stage 16, two clusters were composed of progenitors for more than one trajectory (“bipotent clusters”): a partially biased vagal cluster 9 (*mafb*^+^) resolving into both the cardiac NC cluster 12 and the ENSp NC cluster 13, and a cranial cluster 7 (*nrp2*^+^) which was equally linked to the *rpe65^+^zfhx4^+^* cluster 10 and the *vim^+^itga4^+^* enriched cluster 11 (Figure 2C, Figure 2 – figure supplement 5). From late neural plate stage 14, the unbiased cluster 5, with higher expression of *c9, sox8, tfap2b* in comparison to the earliest unbiased cluster 1, started to split into the vagal-biased 9 (*mafb*^+^, *hoxd3^+^*) and the cranial-biased 7 (*rpe65^+^, dlx2^+^, hox^-^*) trajectories. Despite the global homogeneity within cluster 5, we detected a slight stratification for expression of neuropilins *nrp1* and *nrp2*, which would later mark NC cells emigrating from rhombomere 4 and from rhombomere 2 respectively (Lumb et al., 2014). This suggested that within the unbiased cluster 5, a few cells initiated a trajectory towards cluster 9 (vagal NC, *nrp1, mafb*, *cfi*, *meis2*, *mdk*) while others were shifted towards an anterior hindbrain trajectory (*nrp2*, *id3*, *ifitm3*, *alk*). This indicated that the SC transcriptomes identified an early bias in "bipotent" cell populations, opening hypotheses about gene regulations involved in each fate specification.

Finally, we detected multiple clusters (1, 5 and 14) expressing a canonical NC stem cell-like signature (*sox9, tfap2c, snai2)*, together with high expression of pluripotency markers (*e.g. pou5f1*, *cmyc*, and *sox2),* throughout neurulation. The signature of these potentially multipotent cells, consisted of transcripts common to all NC cells, but was slightly variable across stages (Figure 3D). At, induction stages, this cell lineage expressed enriched levels of *zic1, zic3, tfap2c, cldn6* and *c3*, while at stages 18-22, it was enriched for *tfap2b, mycn, eef1b2, apoc1, vim, rack1, tfap2e* and *c9* expression. Quantitatively, about 72% of NC progenitors were found unbiased at induction stages (gastrula and neural plate stages 12-14), 15% at pre-migratory and EMT stages (stages 16-18) and 9% among tailbud stage 22 NC cells. Interestingly, according to PAGA, the latest unbiased population (14) was transcriptionally equally close to the vagal (9) and cranial (7) transitional clusters, indicating that some stage 18-20-22 cells shared the unbiased pan-NC signature irrespective of their cranial or vagal levels of origin. Moreover, the later unbiased populations (clusters 5, 14) were mostly composed of premigratory cells with low *vim/fn1* expression and remained *hox* negative, generating NC cells which acquired a *hox* signature upon emigration state (*e.g.* hindbrain cluster 15 expressing *hoxa2*).

In sum, this detailed cluster analysis provided the temporal dynamics of gene expression underlying the main transcriptional states defining the various premigratory neural crest trajectories. It highlighted the early specification of several lineages, along with the maintenance of a NC stem-like population at all axial levels between cranial and vagal body axis regions. This analysis further provided a basis for the in-depth exploration of gene program variations which may drive branching at each bifurcation.

**Supplementary Figure Legends**

**Figure 1 - figure supplement 1. A NC classifier based on Xenopus NC selection retrieves NC cells in other whole embryo datasets.**

A binary classifier was trained on the frog NC/non-NC cells (see Methods). Using the whole embryo data matrix of gene expression, we selected the top 10% important features for NC detection. In order to validate it in a whole embryo SC Zebrafish dataset (Wagner et al., 2018), we removed genes present in frog data that were absent in the Zebrafish dataset: Using the subselected top genes, we re-built the model using frog data and obtained the following results: 0.99 accuracy and 0.90 f1 score on a test dataset. When applied to Zebrafish dataset, we obtained also very high scores: the accuracy 0.95 and f1 score 0.66 for 14 stage cells, indicating good initial NC cell selection. **(A)** Top features (gene) for NC classification using LightGBM model. X axis value is the relative importance of the certain gene for the NC classifier **(B)** Confusion matrix indicates the true/false positive and the true/false negative scores for the results obtained on the Zebrafish dataset for stage 14. **(C)** ROC curve for the binary NC classifier indicates high AUC score of 0.95, meaning high recovery of true positives by the model. **(D)** UMAP for Zebrafish stage 14 hpf cells (left) with model-predicted NC cells. The model accurately identifies the cluster of interest (NC-khaki) despite the technical and biological batch effects between Xenopus and Zebrafish datasets.

**Figure 2 - figure supplement 1. Developmental stages and expression of EMT/migration markers per cluster.**

**(A)** Each cluster presents various levels of expression of mesenchymal state/EMT markers and migration regulators. Bar plots with mean normalized expression for each cluster for *vim, twist1, mmp14* and *fn1*. **(B)** Pieplots represent the proportion of cells at each stage in each cluster. Some clusters include cells of 6 successive developmental stages, e.g. cluster 10; while other are almost exclusively found at one single stage.

**Figure 2 – figure supplement 2. Identification of a muscle-like cluster. (A)** At early migration stage (20-22), cluster 16 is strongly enriched for myosin-like genes as well as *bona fide* NC markers, such as *sox10*. **(B, C)** Direct NC induction in ectoderm explants grown *in vitro* (iNC assay) followed by RTqPCR from induction to emigration stages, also identifies the late activation of *myoD* and *myl1* expression at stage 20. This validates the appearence of this muscle-like NC subpopulation among the other NC derivatives already characterized in this NC induction assay (Milet et al., 2013).

**Figure 2 - figure supplement 3. Early-stage biased NC clusters. (A)** From the end of gastrulation, three previously undescribed early trends emerge. Clustering of stage 13 cells only, followed by PAGA analysis identified their similarities with the stage 20-22 clusters: an *olig4*-enriched subpopulation linked to ENSp cluster 13; a *zfhx4-*enriched cluster related to *rpe65*^+^ cluster 10 and a third stage 13 subpopulation expressing *cyp26c1* linked to the cranial NC branch (clusters 4, 6, 15). **(B)** UMAP plots for the early-biased NC subpopulation expressing *hnf1b*, *cdx4*, *nradd* and posterior hox genes. In addition to cluster-specific *hnf1b* and *olig4*, *dhrs3* is co-expressed with *zic5* which specifically stays active during the whole “sacral” trajectory from stage 12 to late ENSp. **(C)** UMAP plots of genes specific for cranial *rpe65+hox-* NC subpopulation specific genes **(D)** UMAP plots of genes specific for rhombomere 4, cranial NC subpopulations.

**Figure 2 - figure supplement 4.** **Early posterior enteric cluster 4 and related late clusters 6, 15. (A)** Line Plot for expression dynamics of *rarg* and *cyp26c1* along trajectories of clusters 4, 6, 15. Cells within clusters are sorted according to pseudotime. **(B)** PAGA plots with *rarg* and *cyp26c1* mean expression in each cluster. The complementary dynamics between levels of expression of the RA-degrading enzyme *cyp26c1* and the RA nuclear receptor *RARg* suggests interesting temporal regulation of retinoic acid-related signaling during development of these NC populations.

**Figure 2 - figure supplement 5.** **Early-stage bipotent NC clusters – cranial vs vagal states.**

UMAP plots for vagal (clusters 9, 12 and 13) and cranial (clusters 7, 10 and 11) specific markers.

**Figure 4 - figure supplement 1.** **TFAP2e connectome generation and validation**

**Table S4, S5. Top-decreased genes after Pax3 or TFAP2e2e depletion.**

Top decreased genes in Pax3 (left) and Tfap2e MO (right) samples with the absolute expression change > 1000.

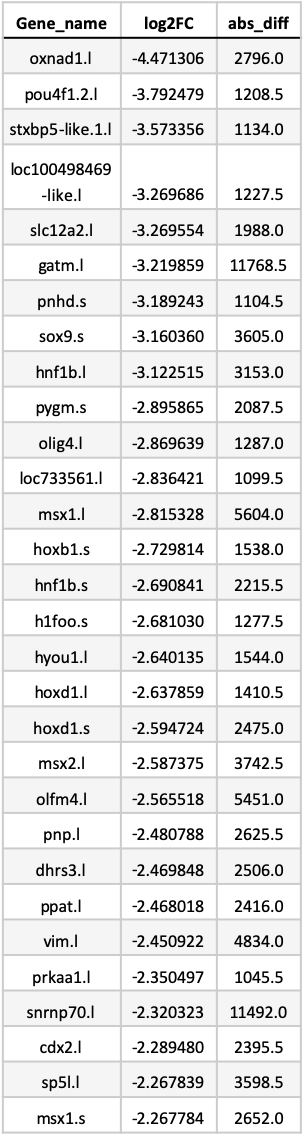

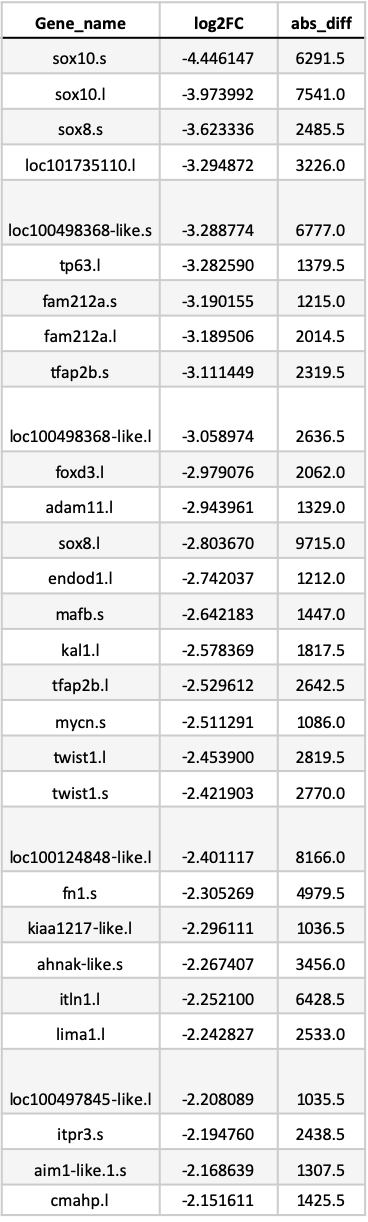

**Table S6. Pax3 targets genes identified by ChIP-seq.**

Top MACS2-scored direct gene targets bound by Pax3.

| **chr** | **start** | **end** | **score** | **gene** |
| --- | --- | --- | --- | --- |
| chr8L | 99475396 | 99475464 | 580 | ASTL3A.1 |
| chr9_10L | 349156 | 349253 | 518 | ATP6V0A1 |
| chr7L | 92931087 | 92931289 | 487 | HES3 |
| Scaffold46 | 813190 | 813276 | 484 | ASS1 |
| chr2L | 176448007 | 176448188 | 448 | PRCP |
| chr8L | 118795321 | 118795461 | 440 | FLAD1 |
| chr8L | 118785517 | 118785631 | 440 | PSMD4 |
| chr7L | 89910794 | 89910942 | 435 | VPS13D |
| chr3L | 130428472 | 130431136 | 417 | S1PR2 |
| chr7L | 15405111 | 15405195 | 390 | INPP5A.1 |
| chr6L | 86407631 | 86407987 | 374 | LYRM4 |
| chr8L | 119048876 | 119048973 | 368 | EFNA3 |
| chr1L | 144729397 | 144731409 | 358 | TBX3 |
| chr5S | 17252239 | 17252405 | 341 | ATF3 |
| chr4L | 141138945 | 141139207 | 341 | ATXN7 |
| chr6L | 37364463 | 37364585 | 314 | HDAC9 |
| chr6L | 124571856 | 124571969 | 312 | ZFAND1 |
| chr6L | 124583418 | 124583619 | 312 | CHMP4C |
| chr4S | 38950018 | 38950121 | 307 | NAE1 |
| chr9_10L | 65487732 | 65488499 | 291 | SLC25A12 |
| chr5L | 51178347 | 51178965 | 288 | PSEN2 |
| chr1S | 115523395 | 115523481 | 286 | NRL |
| chr6L | 39747334 | 39747730 | 286 | EVX1 |
| chr9_10L | 30759439 | 30759550 | 279 | TGM3L.4 |
| chr2L | 88024978 | 88025202 | 267 | HPCA |
| chr6L | 82762756 | 82762905 | 266 | CDH20 |
| chr1S | 12094861 | 12094998 | 263 | FGFR3 |
| chr8S | 33705393 | 33705520 | 248 | FUBP3 |
| chr8L | 57452010 | 57452126 | 245 | SELENOW1 |
| chr9_10S | 23803020 | 23803090 | 244 | USP36 |

**Table S7. TFAP2e ChIP-seq targets.**

Top MACS2-scored genes targets bound by TFAP2e.

| **chr** | **start** | **end** | **score** | **gene** |
| --- | --- | --- | --- | --- |
| Chr7 | 2771023 | 2771259 | 1502 | RBM20 |
| Chr4 | 136928957 | 136929250 | 1360 | PIM1 |
| Chr10 | 10458652 | 10458795 | 1353 | ARL5B |
| Chr10 | 10464454 | 10464497 | 1353 | PLXDC1 |
| Chr5 | 13534757 | 13535087 | 1326 | PTPN14 |
| Chr10 | 50539112 | 50539288 | 1235 | ARPC1A |
| Chr10 | 50529349 | 50529522 | 1235 | PRKAR1A |
| Chr10 | 15734843 | 15734962 | 1218 | SMARCD2 |
| Chr2 | 2687034 | 2687603 | 1212 | BOC |
| Chr9 | 85342807 | 85342887 | 1199 | TNS1 |
| Chr6 | 90349120 | 90349399 | 1198 | TFAP2A |
| Chr6 | 47572683 | 47572824 | 1180 | RARB |
| Chr4 | 58757061 | 58757134 | 1176 | ANKRD11 |
| Chr4 | 48790141 | 48791762 | 1170 | BCAR1 |
| Chr7 | 18235059 | 18235409 | 1164 | JMJD1C |
| Chr2 | 138639744 | 138639788 | 1159 | RBMS2 |
| Chr8 | 6618399 | 6618453 | 1159 | NOTCH1 |
| Chr8 | 142072529 | 142072579 | 1153 | CCT3 |
| Chr8 | 142086526 | 142086803 | 1153 | GLMP |
| Chr2 | 40958194 | 40958575 | 1153 | PDK3 |
| Chr1 | 65662851 | 65663810 | 1145 | SPRY1 |
| Chr4 | 67913743 | 67913832 | 1139 | KCTD15 |
| Chr9 | 83910392 | 83910819 | 1130 | TUBA4A |
| Chr1 | 206833365 | 206833506 | 1100 | RNF165 |
| Chr1 | 16309482 | 16311265 | 1095 | DCTN1 |
| Chr1 | 16305342 | 16305426 | 1095 | ADAM19 |
| Chr10 | 11004405 | 11005055 | 1091 | ARHGAP23 |
| Chr6 | 12882851 | 12882961 | 1084 | DHX30 |
| Chr6 | 12892165 | 12892295 | 1084 | LIMA1 |
| Chr4 | 35478920 | 35479123 | 1071 | PPP1R14B |

**Table S8. PCR** **Primers used in this study.**

| **primer name** | **purpose** | **sequence** | **reference** |
| --- | --- | --- | --- |
| Snail2 Fwd pair1 | ChIP-seq IP quality check | CCATCCCAACACCTGTCGTA | This study |
| Snail2 Rev pair1 | ChIP-seq IP quality check | GCCCACCAGTTCACGTTCAT | This study |
| Snail2 Fwd pair2 | ChIP-seq IP quality check | GCAATGCCTCAGCCTGTGAA | This study |
| Snail2 Rev pair2 | ChIP-seq IP quality check | CAGCGCGTACTGCAATTCATTC | This study |
| Odc Fwd | qRT-PCR | GCCCTTTCTCCCTTTAACGC | (29) |
| Odc Rev | qRT-PCR | TGGTCCCAAGGCTAAAGTTG | (29) |
| Snail2 Fwd | qRT-PCR | CACACGTTACCCTGCGTATG | (77) |
| Snail2 Rev | qRT-PCR | TCTGTCTGCGAATGCTCTGT | (77) |
| Sox10 Fwd | qRT-PCR | CTATTACTGACACACGACGGAGC | (17) |
| Sox10 Rev | qRT-PCR | ACCTCTCATCCTCTGAATCCTGC | (17) |
| MyoD Fwd | qRT-PCR | TACACTGACAGCCCCAATGA | (29) |
| MyoD Rev | qRT-PCR | TGCAGAGGAGAACAGGGACT | (29) |
| Myl1 Fwd | qRT-PCR | GAAACACTTGGGCTGCTTTCTT | This study |
| Myl1 Rev | qRT-PCR | AGCAGGTTTAGCCTCAGGTTT | This study |

**Supplementary Materials and Methods**

***Experimental Design***

Single cell transcriptomes from developing frog embryos were scrutinized for neural crest progenitor development using machine-learning tools to infer the gene regulatory network connectome and the gene programs underlying branching of fates. These predictions were largely validated *in vivo* using micro-manipulations in *X. laevis* embryos followed by RNA-seq or ChIPseq.

***Single cell sequencing***

No new materials were collected for this study. Instead, we re-sequenced the RNA libraries for developmental stages NF11 to NF22 (Faber and Nieuwkoop, 2020) used in (Briggs et al., 2018). Please refer to the methods therein for staging, embryo collection, dissociation, cell collection, barcoding (inDrop v2 and v3) and library preparation methods. For re-sequencing, we used two flow cells (two lanes each) of NovaSeq S2 at 100 cycles setting generating a total of 12 billion reads. All datasets are deposited under NCBI Gene Expression Omnibus number GSE198494.

***Reference genome and gene symbol assignments***

For bioinformatics analysis of SC dataset, we used the *X.tropicalis* v.10 genome assembly, gene models v. 10.7. Specifically, we used the file Xentr10-Gene-Sym-HUMAN-BLOSUM45.txt downloaded from Xenbase on Nov 5th, 2021. Thousands of genes in that transcriptome version lack interpretable gene symbols, though in many cases an unambiguous protein identity can be identified by sequence homology. To fill the holes in gene symbol assignments we assigned protein gene symbols to each transcript using a modified reciprocal best HMMER hit approach (Savova et al., 2017) based on a target reference set of curated human proteins.

***scRNA-seq reads processing***

Compared to (Briggs et al., 2018), we selected a different aligner and scRNA pipeline since Bowtie (Langmead et al., 2009), which was used in the InDrops pipeline (Klein et al., 2015), was not designed to align transcripts to the genome, and therefore is not splice-aware. A splice-aware algorithm does not align RNA-seq reads to introns, and identifies possible downstream exons trying to align to those instead. Moreover, using Bowtie for aligning reads to a transcriptome raises questions about choosing the correct transcript. Therefore, we chose STAR, since it is one of the best RNA-seq aligners and outperforms other aligners in terms of correctly and incorrectly aligned reads (Baruzzo et al., 2017). Also, it is fast and there is no need to do preprocessing by removing bad quality bases or adapters as STAR does that internally (Dobin et al., 2013). We used the DropEst pipeline (Petukhov et al., 2018). Recent benchmarking shows that DropEst exceeds other options in terms of sensitivity and efficiency (running time and memory use) (Gao et al., 2021). As a result, for v3 we used Indrops demultiplexing and soft filtering, then we generated tagged files similar to dropTag without considering reads base quality for barcodes, followed by alignment and quantification with STAR (outFilterMatchNminOverLread 0.44 --outFilterScoreMinOverLread 0.44) and dropEst using X.trop v10. After filtration by counts and genes numbers (>200 genes; >300 counts), we gathered a dataset of 177250 cells. In the cells of interest (Ectoderm and NC cells), mean counts number was 1778, and mean gene number was 1035.

***scRNA-seq postprocessing***

To process scRNA data we used Scanpy (Wolf et al., 2018), a comprehensive scRNA pipeline that is functionally similar to Seurat (Stuart et al., 2019). Firstly, we calculated the percentage of mitochondrial (MT) and ribosomal (RB) genes per cell,by manually calculating the fraction of mitochondrial reads and ribosomal reads. As high MT proportions indicate poor quality cells (Lun et al., 2016), presumably due to loss of cytoplasmic RNA from perforated cells, we removed cells with a high percentage of MT/RB transcripts. Then, we removed gene sets which might affect the normalization step. These include foreign tissue contaminants, heatshock, ribosomal, clock and hemoglobin genes. For normalization, we excluded the expression of genes if their expression was more than 3 percent of the total expression of the cell, because it could greatly affect the resulting normalized values for all other genes (Weinreb et al., 2018). Next we assessed the cell cycle dynamics. The algorithm calculates the difference between the mean of the given list of cell cycle genes and the mean of the reference genes. To build such a reference, we randomly selected a set of genes that matched the distribution of the expression in the list. This way we got S phase, G2M phase scores for each cell. We did not regress out cell cycle effect because it had no significant influence. For normalization, we used scanpy function *scanpy.pp.normilize_total* without highly expressing genes and target_sum=10^4.

***Clustering and NC cells selection***

For each stage, we performed independent standard dimensionality reduction with PCA, computing a neighborhood graph (n_pcs selected based on the amount of variance explained by each PC, n_neighbors selected manually) and UMAP (Jacomy et al., 2014). For clustering, we used the Leiden algorithm (Traag et al., 2019). For each cluster, we defined cluster-specific genes with differential expression analysis (scanpy t-test_overestim_var) and selected only clusters which were the most similar to NC cells using NC signatures from (Briggs et al., 2018). For NC clustering, we defined the optimal number of clusters by manually increasing their number and checking for biological meaning, as revealed by specific gene expression, for example of *hox* genes, such as *hoxd3* for cardiac NC (cluster #12), or *hoxb6* for Enteric Nervous System progenitors (ENSp, cluster #13).

***Binary NC classifier***

First, we generated a dataset by labeling cells as NC based on the previous differential expression analysis between clusters (Briggs et al., 2018). For feature selection, we trained the LightGBM model (Ke et al., 2017) on the whole embryo dataset of 177250 cells, retrieved the top 2000 important features and used it for re-training the model (default setting with n_estimators=500). The resulting model detected NC cells in the test dataset from the *Xenopus tropicalis* dataset with accuracy 0.99 and F1 score 0.90. Only the trained dataset sample was used to get the top important features and retrain the model on the top 2000 features. To test the model robustness, we validated the NC classifier in the Zebrafish scRNA-seq dataset (Wagner et al., 2018). We used a sigmoid function to smoothen the technical batch effect between the datasets. Genes that were not found in the Zebrafish dataset were replaced with NaN values (728 from top 2000 important genes were not found in the Zebrafish dataset). Finally, our model predicted NC cells in the Zebrafish dataset with AUC score 0.95 and F1 score 0.66 (in comparison, the random model with strategy="stratified" (https://scikit-learn.org/stable/modules/generated/sklearn.dummy.DummyClassifier.html) applied on the imbalanced Wagner dataset has F1 score=0.05). This result is striking because it was robust despite significant species-specific variations in the expression of classical NC gene markers between frog and fish, and despite a strong batch effect between the datasets (595 genes from the list of the most important genes for NC classification were missing from the zebrafish dataset). For example, the expression of *c9* and *snai2,* which are pan-NC genes in *Xenopus,* was drastically different in the zebrafish NC dataset. Yet, the classifier efficiently recognized NC cells in the fish neurula. This result confirmed the accuracy and robustness of our NC cells selection criteria.

***GRN generation***

Using the temporal dynamics of gene expression, several algorithms have been designed to infer genetic co-regulation (Pratapa et al., 2020). One of the best tools, GRNBoost2, is based on a gradient boosting machine and retrieves the gene regulatory network (GRN) from the expression data-matrix (Moerman et al., 2019). Starting from a given list of TFs, this algorithm powerfully links genes with similar expression patterns in SC transcriptomes, providing “syn-expression groups” (Niehrs and Pollet, 1999) at the single cell level. Thus, GRNBoost2 provides a network of potential bidirectional gene relationships (including genes encoding TFs and other types of proteins). Through this algorithm, for each gene in the cell-gene matrix, a tree-based regression model is built to predict gene expression using the expression of TFs. Each gene-specific model produces a partial GRN with regulatory associations from the most predictive TFs for the gene. Then, all regulatory associations are pooled and ranked by importance to complete the GRN. The *Xenopus* TF list (1417 TFs) was taken from *Xenopus tropicalis* TF catalog (Blitz et al., 2017). We analyzed the resulting network of TFs and their targets using the networkx package and identified the most important nodes by calculating betweenness and degree centralities with betweenness_centrality, degree_centrality network functions.

***Principal graph generation***

The tree analysis was carried out using the scFates package (Albergante et al., 2020), based on ElPiGraph using the concept of elastic energy and a gradient descent-like optimization of the graph topology (Albergante et al., 2020). However, this approach is too sensitive to build the principal graph for the whole NC dataset. Therefore, to generate the main tree, we used the PAGA algorithm (Wolf et al., 2018). This revealed cluster-cluster relationships including the early stages where the strongest connectivity was observed. Further we used ElPiGraph to study specific branches and bifurcation points.

***Branching analysis***

Using ScFates we defined features significantly changing along the tree, and then using pseudotime values and differential expression analysis, determined early and late branch-specific features. For each bifurcation point of interest, we selected a set of cells related to the clusters involved in the bifurcation. The selection of parameters for building the principal tree for each point of the bifurcation was carried out using brute force approach.

***Chromatin immunoprecipitation sequencing (ChIPseq)***

Chromatin immunoprecipitation was performed according to (Wills et al., 2014). Embryos were injected in both blastomeres at the two-cell stage with tracing amounts (75 pg) of mRNA encoding either Pax3-FLAG-HA, or TFAP2e-FLAG. Injected embryos were collected at mid-neurula stage 14 (Pax3, 100 *X. laevis* embryos/condition, three biological replicates) or at NC early migratory stage 19 (TFAP2e, 100 *X. tropicalis* embryos/condition, one biological replicate). IP efficiency was tested on *snai2* promoter (Table S8). After sequencing, 100 bp single-end reads were aligned to *X. laevis* genome version 9.2 or *X. tropicalis* v10.0 using Bowtie2. Peaks were called using MACS2. For Pax3 and TFAP2a, we selected peaks common in three replicates, for TFAP2e we used stricter MACS2 score cutoff=500. Target genes were searched with bedtools (window size = 10kb).

***In vivo experiments: Xenopus laevis injections, microdissections, grafting and small RNA-seq***

Animal care and use for this study were performed in accordance with the recommendations of the European Community (2010/63/UE) for the care and use of laboratory animals. Experimental procedures were specifically approved by the ethics committee of the Institut Curie CEEA-IC #118 **(Authorization APAFiS#36928-2022042212033387-v1 given by National Authority)** in compliance with the international guidelines.

*In vivo* injections, NB/NC dissections and grafting were done as previously (Milet and Monsoro-Burq, 2014; Plouhinec et al., 2017) using *X. laevis* embryos. For knockdown experiments, previously validated antisense morpholino oligonucleotides (MO) were used to deplete *pax3,* and *TFAP2e* transcripts (GeneTools). Pax3 MO (20ng) (Monsoro-Burq et al., 2005) or *TFAP2e* MO (20ng) (Hong et al., 2014). Depletion efficiency was checked using *in situ hybridization* on sibling embryos to verify reduction of *snai2* expression (not shown). One Pax3 morphant anterior NB explant (stage 14) or one TFAP2e-morphant NC explant (stage 17) were dissected from the injected side, in biological triplicates. After RNA extraction and cDNA library preparation, each individual explant was sequenced (small RNA-seq). The resulting 100 bp paired-end sequencing reads were aligned to the *X. laevis* genome version 9.2 using STAR and the count reads were analyzed using String Tie. Differentially expressed genes were selected considering log2FC and expression difference in absolute values (abs. diff. >=100 and <=500: log2FC>1.5 or log2FC<-1.5; abs. diff >=500 and <=1000: log2FC>1 or log2FC<-1; abs. diff >=1000 and <=3000: log2FC>0.5 or log2FC<-0.5; abs.diff.>3000: log2FC>0.33 or log2FC<-0.33)

***iNC assay, RNA quantification and RT-qPCR***

The induced neural crest assay (iNC) used co-activation of dexamethasone-inducible Pax3-GR and Zic1-GR at gastrula initiation stage 10.5, in pluripotent blastula ectoderm (animal caps dissected at blastula stage 9) (Hong and Saint-Jeannet, 2007; Milet et al., 2013). This was combined with Sox9 depletion (40 ng of *sox9* MO) (Spokony et al., 2002), or gain-of-function (300 pg *sox9* mRNA). At the desired stage, 8-10 explants/condition were harvested and processed for RTqPCR, in biological duplicates, as in (Alkobtawi et al., 2021). Primers listed in Table S9.

***Whole-mount in situ hybridization (ISH)***

Whole-mount *in situ* hybridization followed an optimized protocol (Monsoro-Burq, 2007). Embryos were imaged using a Lumar V12 Binocular microscope equipped with bright field and color cameras (Zeiss).

***Benchmarking SC-based machine-learning and in vivo experimental approaches to build the neural crest GRN.***

Our SC analyses provide a genome-wide connectome based on a total of 16978 interactions computed between 405 TFs (out of the initial list of 1417 TFs expressed in developing embryos) and 4532 other genes expressed in neural crest during gastrulation and neurulation. The biological significance of the network was validated using experimental analysis for three different NB/NC specifiers, Pax3 and TFAP2e, as transcriptomic interactions predicted using GRNBoost2 retrieved many interrelations supported either by direct TF binding (ChIP) or by TF depletion *in vivo*. However, when comparing the targets predicted with SC transcriptome modelling and experimental validation *in vivo*, we find that the fraction of gene correlations predicted with GRNboost2 also validated by ChIPseq was 7-9%, and also confirmed by TF depletion (which includes both direct and indirect targets) was 11-16%. Changing the network filtration criteria (weights, expression level) did not affect the proportion of validated genes. This means that despite the significant discovery power displayed by GRNBoost2 on a complex dataset, there is margin for increasing its specificity and accuracy. For example, GRNBoost2 did not link *tfap2e* to *hnf1b* (although confirmed by both ChIPseq and MO depletion), possibly because those genes have quite different expression patterns during the course of neurulation (Figure 4 - figure supplement 2B). Likewise, GRNBoost2 did not link *tfap2e* to *notch1* either (also confirmed with ChIPseq and MO) although they display closely related patterns. Another hypothesis for the incomplete coincidence of the predictions between MO/ChIP and GRNboost2 is that the dataset compiled for GRNboost2 includes cells of all available stages (due to the need for a large number of cells for modelling) while the MO/ChIP experiments were carried out at specific stages. Additionally, another limitation of GRNboost2 is its weak ability to predict the relationship between genes with different expression time-frames, i.e. when a gene A activates a gene B and stops being expressed in B-expressing cells. To compare these results to a random chance we utilized bootstrapping. To do this, we randomly selected the same number of genes for different methods (ChIP-seq, MO, GRNboost2) 1000 times from the total number of genes (*X.laevis* and *X.tropicalis*). We obtained a median intersection value of 3, confirming that our analyses were not obtained at random.
